# Bacterial tolerance to host-exuded specialized metabolites structures the maize root microbiome

**DOI:** 10.1101/2023.06.16.545238

**Authors:** Lisa Thoenen, Caitlin Giroud, Marco Kreuzer, Jan Waelchli, Valentin Gfeller, Gabriel Deslandes-Hérold, Pierre Mateo, Christelle A.M. Robert, Christian H. Ahrens, Ignacio Rubio-Somoza, Rémy Bruggmann, Matthias Erb, Klaus Schlaeppi

**Affiliations:** Institute of Plant Sciences, University of Bern, 3013 Bern, Switzerland; Department of Environmental Sciences, University of Basel, 4056 Basel, Switzerland; Interfaculty Bioinformatics Unit, University of Bern, 3012 Bern, Switzerland; Agroscope, Molecular Ecology and Swiss Institute of Bioinformatics, Zurich, 8046 Zürich, Switzerland; Molecular Reprogramming and Evolution (MoRE) Lab, Centre for Research in Agricultural Genomics (CRAG), Carrer Vall Moronta Edifici CRAG, 08193, Barcelona, Spain

**Author notes:** **Corresponding author:** Klaus Schlaeppi. **Author Contributions:** L.T., M.E., and K.S. designed research; L.T. and M.K., performed research, J.W. supported bioinformatic analysis, C.G, C.S., V.G., G.D.-H. supported experiments, P.M. and C.A.M.R. provided experimental reagents and materials and supported experiments, C.H.A., I.R.-S., and R.B. contributed to genome sequencing, provided new analytic tools, L.T., M.K, M.E., and K.S. analyzed data and L.T., M.E., and K.S. wrote the first draft of the paper. All authors revised the paper. **Competing Interest Statement:** The authors declare no competing interest.

## Abstract

Plants exude specialized metabolites from their roots and these compounds are known to structure the root microbiome. However, the underlying mechanisms are poorly understood. We established a representative collection of maize root bacteria and tested their tolerance against benzoxazinoids, the dominant specialized and bioactive metabolites in the root exudates of maize plants. *In vitro* experiments revealed that benzoxazinoids inhibited bacterial growth in a strain- and compound-dependent manner. Tolerance against these selective antimicrobial compounds depended on bacterial cell wall structure. Further, we found that native root bacteria isolated from maize tolerated the benzoxazinoids better compared to non-host Arabidopsis bacteria. This finding suggests the adaptation of the root bacteria to the specialized metabolites of their host plant. Bacterial tolerance to 6-methoxy-benzoxazolin-2-one (MBOA), the most abundant and selective antimicrobial metabolite in the maize rhizosphere, correlated significantly with the abundance of these bacteria on benzoxazinoid-exuding maize roots. Thus, strain-dependent tolerance to benzoxazinoids largely explained the abundance pattern of bacteria on maize roots. Abundant bacteria generally tolerated MBOA, while low abundant root microbiome members were sensitive to this compound. Our findings reveal that tolerance to plant specialized metabolites is an important competence determinant for root colonization. We propose that bacterial tolerance to plant-secreted antimicrobial compounds is an underlying mechanism determining the structure of host-specific microbial communities.

**Significance Statement:** Diverse microbial communities colonize plant roots. They feed on carbon rich root exudates which contain a diverse mix of chemicals including primary and specialized metabolites. Here we show that specialized metabolites act as selective antibiotics to shape the root bacterial communities. By growing single isolates of maize root bacteria in the presence of benzoxazinoids *in vitro*, we find that the strains differ greatly in their tolerance to benzoxazinoids. Their different levels of tolerance largely explained their abundance on benzoxazinoid-exuding roots. Our work shows how plant specialized metabolites act to shape the maize root microbial community and thus deepened our mechanistic understanding of how plants shape their microbiome.

## Introduction

Diverse communities of microbes including bacteria, fungi, oomycetes, and protists colonize plant roots (1–3). Collectively functioning as a microbiome, these microbes provide several benefits to their host. They can improve plant growth through the production of plant hormones (4, 5), improve plant nutrient uptake (6–8), and protect plants against pathogens (9–11). The root microbiome is mainly recruited from soil and thus its composition resembles the surrounding soil microbiome (1, 12). Plants further shape the composition of their root microbiome by root morphological traits (13, 14), immune responses (15, 16), and secretion of diverse root exudates (17–19). Through root exudation, plants release up to 25% of their assimilated carbon to the surrounding soil (20, 21). Soil microbes primarily exploit the exudates as carbon substrates for growth and are therefore attracted to the roots (22, 23). Such primary carbon substrates include sugars, amino acids, or organic acids, but importantly, root exudates also contain plant specialized metabolites (17, 18, 24).

Plant specialized metabolites govern the interactions of plants with the environment (25). Among numerous functions, they were shown to shape the root and rhizosphere microbiomes (17, 18, 26). Of note, microbiome structure is not solely defined by plant-but also by microbe-derived specialized exometabolites (27). Studies on plant metabolites often compare microbiome composition on roots or rhizospheres of wild-type plants relative to biosynthesis mutants defective in the production of a specialized metabolite. Glucosinolates (28), camalexin (29), triterpenes (22), and coumarins (30–32) were shown to structure the root microbiome of the model plant *Arabidopsis thaliana*. Sorgoleone exuded by sorghum (*Sorghum bicolor*) (33), gramine produced by barley (*Hordeum vulgare*) (34) and benzoxazinoids (35–38), diterpenoids (39), zealexins (40) and flavonoids (14) released by maize (*Zea mays*) shape the root microbial communities of their respective hosts. While it is well documented that plant specialized metabolites present drivers for root microbiome assembly, the underlying mechanisms that explain community structure remain largely unknown.

Benzoxazinoids (BXs) are specialized compounds produced by sweet grasses (*Poaceae),* which include important crops such as maize, wheat, and rye (41). These indole-derived alkaloids are especially abundant in young maize seedlings and actively growing tissues, accounting for up to 1% of plant dry weight (42). Besides structuring the root and rhizosphere microbiomes (35–38), BXs defend against insect pests and pathogens and play a role in defense signaling (43). Further, BXs function as phytosiderophores by chelating iron and thus improving plant nutrition (44). The major BX exuded by maize to the surrounding rhizosphere is DIMBOA-Glc (26; see Table S1 for full names and structures of BX compounds). In the rhizosphere, it is deglycosylated to DIMBOA and rapidly converted to MBOA, where it is stable for days to weeks (45). Through microbial activity, MBOA is further converted to AMPO, which then accumulates and remains detectable in the soil for months to years (45–47). AMPO, an aminophenoxazinone, has allelopathic function by suppressing the growth of neighboring plants (45, 48–50). While maize primarily synthesizes methoxylated BXs, rye or barley mainly produce non-methoxylated analogs following the same chemical conversion pathway (DIBOA-Glc > DIBOA > BOA > APO, see Table S1; 41, 48, 56). To what extent different BX compounds contribute to microbiome structuring remains to be investigated.

BXs are known to affect microbes. Diverse responses, mainly of individual bacterial strains, have been reported for various BX compounds. On one end, BXs are toxic as demonstrated with pure compounds such as MBOA that inhibited the growth of bacteria like *Streptococcus aureus* and *Escherichia coli* (52). On the other end, DIMBOA-derived compounds affect bacterial behavior. They were shown to reduce bacterial motility and biofilm formation of the bacterial pathogen *Ralstonia solanacearum* (53). DIMBOA was shown to act as a chemoattractant for host localization by the beneficial rhizobacterium *Pseudomonas putida* (54). HDMBOA, a dimethoxylated form of DIMBOA, was found to reduce the virulence of the pathogen *Agrobacterium tumefaciens* (55). A few studies have addressed how bacteria cope with BX compounds. For instance, strains of *Pantoea* and *Bacillus* species can degrade BXs and convert them into various metabolites (56, 57). Some other bacteria were found to tolerate BXs, such as e.g. *Photorhabdus,* an endosymbiotic bacterium of nematodes, that tolerates high levels of MBOA (58). Mechanistically, this tolerance to MBOA was based on an aquaporin-like membrane channel *aqpZ*. The antimicrobial activity of BXs strongly differs among taxonomically diverse bacteria, as shown for Arabidopsis root strains when testing their growth in the presence of the non-methoxylated BX compounds BOA and APO (59). While this study convincingly revealed strain-level antimicrobial activity of BXs, it remains unclear whether bacterial tolerance to plant specialized metabolites could explain the structure of root microbiomes.

Given that BXs generally structure the microbiome of maize roots (35–38), that exudation of BX leads to accumulation of MBOA in the rhizosphere (37), and that bacteria widely differ in their tolerance to BXs (59), we hypothesized that bacterial tolerance to BXs, in particular to MBOA, explains the BX-dependent community structure of the maize root microbiome. To test this hypothesis, we established a strain collection from roots of BX-exuding maize plants, similar to culture collections of other plant species including Arabidopsis (60), clover (61), rice (62), lotus (63), and maize (64, 65). Strain collections, which bridge cultivation with culture-independent sequencing methodologies, present powerful resources for the in-depth studying of the molecular mechanisms in microbiomes (66). In a second step, we screened the maize root bacteria for their tolerance against pure BXs and aminophenoxazinones present in the maize rhizosphere. Among the 52 isolates tested, we found a broad spectrum of strain-level tolerances to MBOA without a phylogenetic clustering. Mapping these strains to root microbiome profiles revealed that their levels of tolerance to MBOA largely explained their BX-dependent abundance on maize roots. Overall, our findings reveal that a bacterium’s tolerance to the root-secreted antimicrobials is a key trait for defining its abundance in the root microbiome.

## Results

### Benzoxazinoid exudation does not alter bacterial community size on maize roots

Given their antimicrobial activities, we hypothesized that BXs may reduce the overall community size which represents the total number of microbial cells. Thus, we quantified bacterial community size (i) by plating root extracts and scoring colony-forming units and (ii) by quantitative PCR measurements of bacterial DNA relative to plant root DNA. We compared roots of BX-exuding (wild-type) and BX-deficient (*bx1* mutant) maize in two greenhouse experiments with field soil. Additionally, we quantified bacterial community size using DNA extracts of earlier field experiments with wild-type and *bx1* plants (35). For the greenhouse experiments, the cultivation-dependent quantifications of bacterial cell numbers and the load of bacterial DNA on roots did not differ between the two genotypes; findings that were further confirmed by the field samples (Fig. S1). Thus, we conclude that BX exudation, at least under the conditions tested, does not affect the community size of the bacterial maize root microbiome.

### Culture collection covers abundant bacteria of the maize root microbiome

We built a culture collection of maize root bacteria (hereafter, MRB collection) isolated from roots of wild-type B73 maize grown in natural soil in the greenhouse. We used the same soil from the Changins site (CH) where the structuring of the maize root microbiome by BXs was first observed (37). We determined the taxonomy of each isolate by sequencing parts of the 16S rRNA gene using Sanger technology. The MRB collection consists of 151 bacterial isolates representing 17 taxonomic families across the five major phyla Pseudomonadota (n = 69), Actinobacteriota (n = 56), Bacillota (n = 23), Bacteroidota (n = 2) and Deinococcota (n = 1; Fig. 1). Among those were typical root-colonizing families such as Pseudomonadaceae, Microbacteriaceae, Rhizobiaceae and Sphingomonadaceae (1). To investigate the abundance of community members (ASV / OTU) corresponding to MRB strains in profiles of maize roots from, we mapped the 16S rRNA Sanger sequences to microbiome profiles of the roots, from which they were isolated (37). This analysis revealed that MRB strains mapped to taxonomic units that accounted for 24% of the total microbiome abundance at the sequence level (Fig. S2A) with 112 isolates being abundant (>0.1% abundance, respectively) and 34 low abundant (<0.1%) community members (Fig. 1, inner ring).

**Figure 1:**
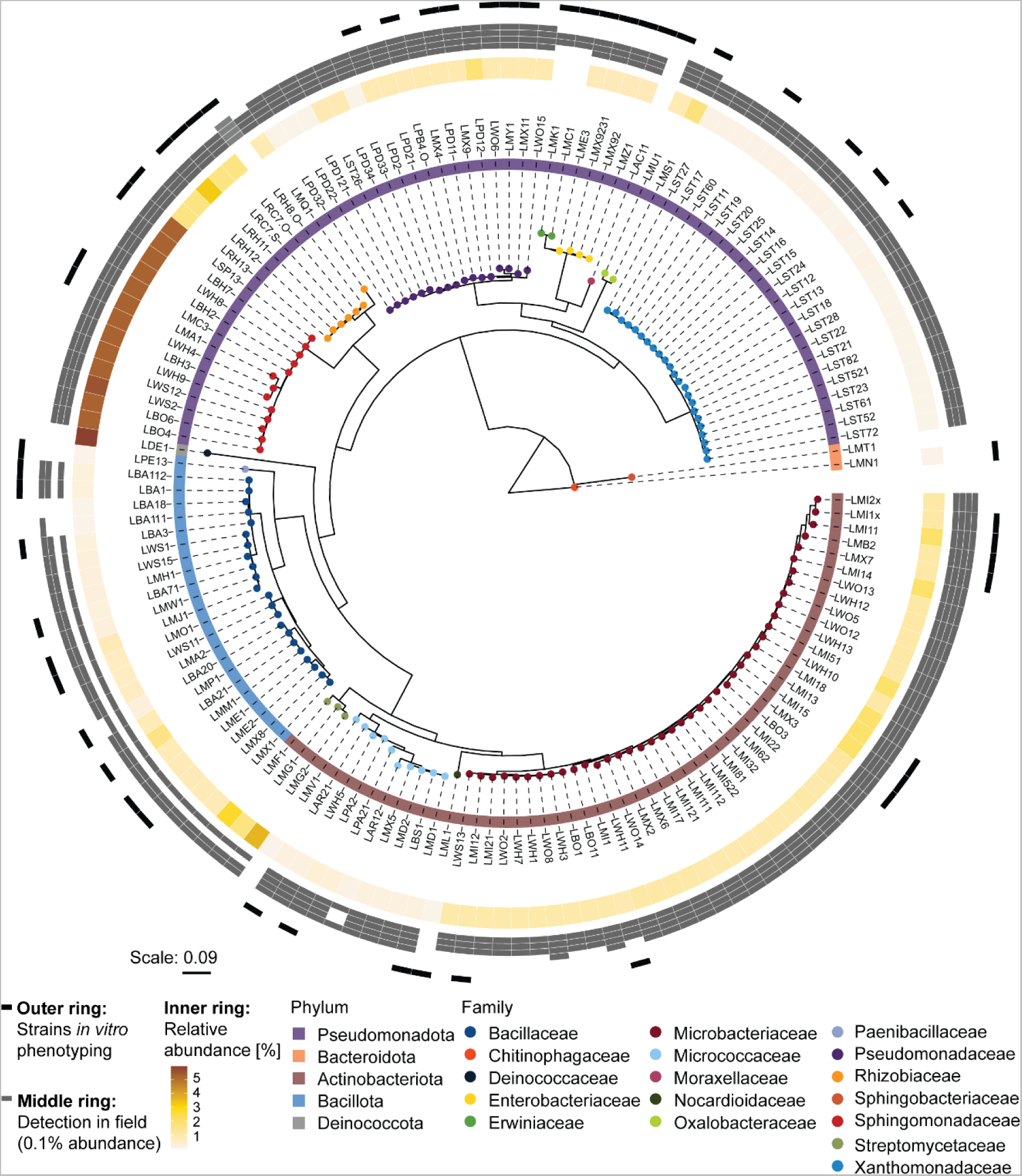
Maize root bacteria (MRB) strain collection. Maximum likelihood phylogeny, constructed from the alignment of 16S rRNA gene sequences. Leaf nodes are colored by family taxonomy and the ring next to the strain IDs reports phylum taxonomy. The rings on the outside of the tree show the abundance of the strains in the microbiome datasets: The inner ring is colored according to the relative abundance (%) of the corresponding sequence in the microbiome profile of the roots, from which the isolates were isolated from (origin of isolation). Grey boxes in the middle ring mark when a strain was detected (> 0.1 % abundance) in root microbiome profiles of maize grown in field soil (Changins – CH, Reckenholz – CH, Aurora – US, and Sheffield – UK). The outer ring denotes in black the representative strains used for growth assays.

Next, we assessed whether the MRB isolates could be detected in root microbiomes of maize grown in the field and other soils. We mapped the 16s rRNA sequences of the MRB isolates to the following microbiome datasets: field-grown maize in Changins (26), Reckenholz (both CH), and Aurora in the US (35), and a pot experiment with field soil from a location in Sheffield, UK (36) (Fig. 1, middle ring). Consistent with the pot experiments for isolation, the majority of MRB isolates (139/151) also mapped to abundant (>0.1%) root microbiome members of maize grown in the field in Changins from where the soil was used for the isolation experiments (Fig. S2B). Similarly, most MRB isolates were also detected as abundant members in maize root microbiomes in other field soils from Switzerland (117/151), the US (140/151), and the UK (84/151). Isolates belonging to Pseudomonadaceae, Microbacteriaceae, and Oxalobacteraceae were detected in all four soils as abundant members. To make the MRB strain collection a useful resource for further studies, we sequenced the genomes of selected strains (Dataset S1). Taken together, these results demonstrate that the MRB collection taxonomically covers the major bacterial families and largely represents the abundant members of maize root microbiomes in different field soils.

### MBOA is the most selective exudate compound to inhibit maize root bacteria in vitro

To investigate the tolerance of MRB strains to BXs, we screened a representative set of 52 isolates (Fig. 1, outer ring) using *in vitro* growth assays with purified DIMBOA-Glc as well as synthetic MBOA and AMPO. We did not include DIMBOA, the bioactive aglycon of DIMBOA-Glc (37), as it was immediately converted to MBOA in our screening system (data not shown). We exposed the MRB strains to varying compound concentrations and measured bacterial growth based on optical density in liquid cultures over time. We calculated the area under the bacterial growth curve (AUC), normalized it to the growth in the control treatment, and defined a tolerance index (TI) for each strain across all tested concentrations (See methods and Fig. S3). In contrast to defining medium inhibitory concentrations (IC_50_), the TI method allows tolerance comparisons among strains including ones that are not inhibited at the highest tested concentration. The maximal TI value 1 indicates that the strain is not inhibited. We defined bacteria with TI values 0.75-1 as *tolerant*, with values 0.5–0.75 as strains of *intermediate tolerance*, and values <0.5 classify *susceptible* strains. Fig. S3 exemplifies the approach with the MBOA-tolerant *Pseudomonas* LPD2 (inhibited only at the highest concentrations, TI = 0.88) and the MBOA-susceptible *Rhizobium* LRC7.O (inhibited already at the lowest concentration, TI = 0.25).

Figure 2 reports the growth of all tested MRB strains to DIMBOA-Glc and TI to MBOA and AMPO (growth in MBOA and AMPO, see Figs. S4 & S5). For DIMBOA-Glc, the main compound secreted by maize roots (Table S1), we could only test two concentrations due to the limited availability (500 and 2500 µM). The majority of MRB strains was tolerant to 2500 µM (AUC > 0.75, 43/52 strains) with some strains even benefiting with improved growth (AUC > 1, 21/52; Figs. 2A & S4). DIMBOA-Glc inhibited the growth of only six strains, predominantly belonging to the family of Bacillaceae.

**Figure 2:**
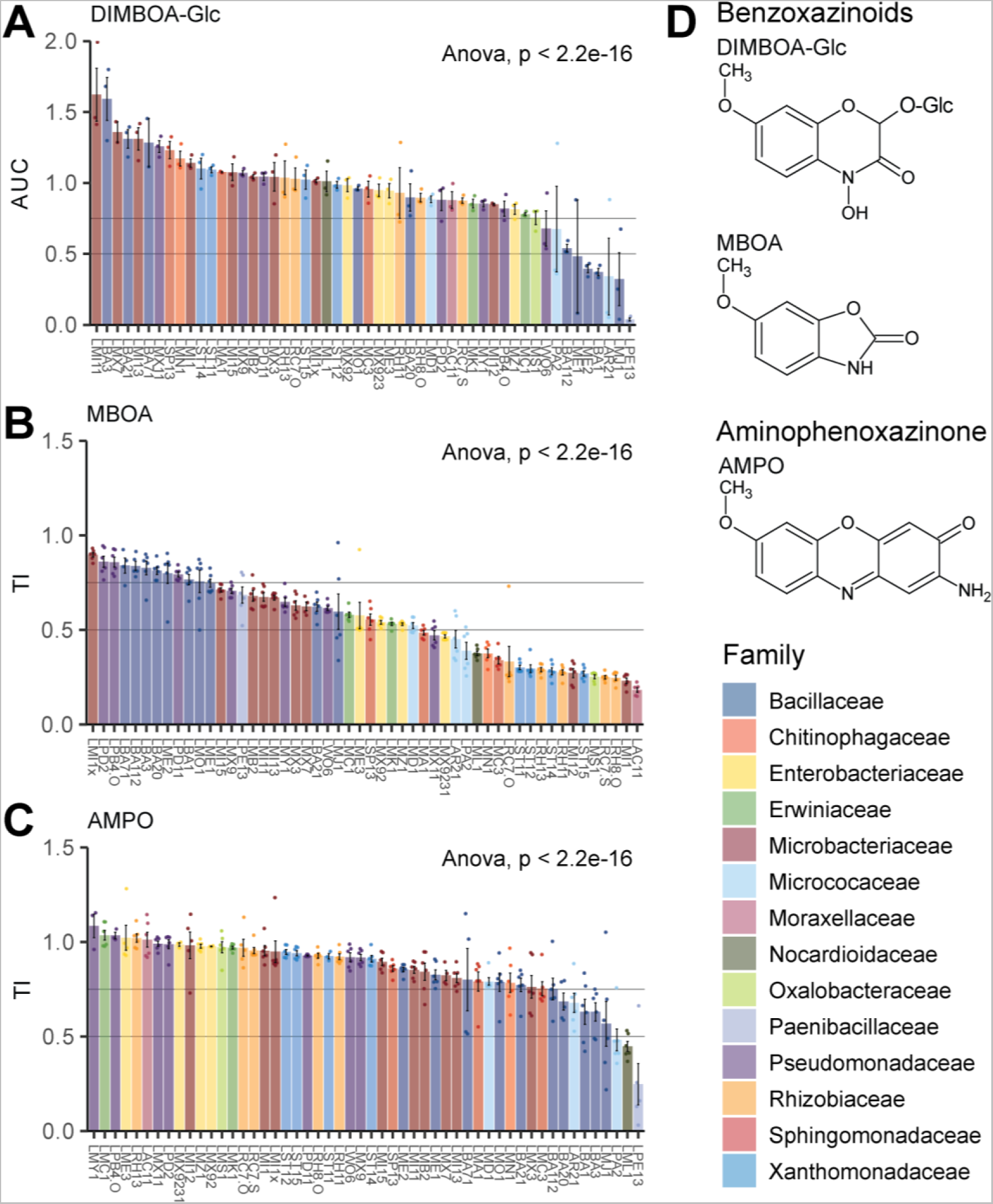
Taxa- and compound-specific growth responses of maize root bacteria to benzoxazinoids and aminophenoxazinones. **A)** Bacterial isolates are colored according to family taxonomy and their growth (AUC value in 2500 μM) in DIMBOA-Glc and their tolerance indices (TI) for **B)** MBOA and **C)** AMPO are shown. Bar graphs (A-C) showing means ± SE bar with individual data points (n = 3 for DIMBOA-Glc, n = 6 for MBOA and AMPO). Results of ANOVA (model: TI ∼ strain) are shown inside the panels.

In contrast to DIMBOA-Glc, MBOA, which is the major compound detected in the rhizosphere (Table S1) and known for its antimicrobial activity (52, 54, 58), strongly inhibited more than a third of the strains (22/52 strains) and moderately affected another third of the MRB strains (18/52 strains; Fig. 2B & S5A). The most susceptible strains belonged to the Rhizobiaceae and the Moraxellaceae families. Only 12 strains were tolerant to MBOA (TI > 0.75), belonging to Pseudomonadaceae, Bacillaceae, and Microbacteriaceae. Strains belonging to the same family typically showed a similar tolerance level to MBOA (Fisher exact test: tolerance group ∼ family, p < 0.001). Among Microbacteriaceae we found the strongest phenotypic heterogeneity, ranging from the most tolerant strain LMI1x to the second most susceptible strain LMI1 and with many strains of intermediate tolerance.

Finally, we tested aminophenoxazinone AMPO, the direct microbial degradation product of MBOA that accumulates at low levels in the rhizosphere (Table S1). Because AMPO is insoluble at concentrations > 50 µM, it was not tested up to the same concentrations as DIMBOA-Glc and MBOA. A separate experiment directly comparing bacterial growth in 50 µM of MBOA vs AMPO revealed that the highest testable concentration of AMPO inhibited growth of several MBOA tolerant strains (Fig. S6A). This suggested that concentrations up to 50 µM of AMPO would cover the necessary dynamic range of its toxicity. Most MRB strains were tolerant (43/52 strains) or only moderately affected (6/52 strains), and only 3 strains were susceptible to AMPO (TI < 0.5; Fig. 2C & S5B). The affected strains belonged to Bacillaceae and Micrococcaceae (Fisher exact test: tolerance group ∼ family: p < 0.05). Comparing bacterial tolerances of AMPO with MBOA revealed a weak, yet significant negative correlation, meaning that AMPO-tolerant bacteria were not necessarily also MBOA-tolerant

(Fig. S5C). The direct comparison of toxicities (analyzing bacterial growth in AMPO vs. MBOA, both at 50 µM; Fig. S6B) revealed that only one strain was significantly affected in growth by 50 µM MBOA (LWO6; T-test: p < 0.05) while twenty strains showed growth reduction by the same concentration of AMPO (T-tests: p < 0.05). This reveals that AMPO is more toxic than MBOA. However, AMPO only accumulates at very low levels in the rhizosphere (Table S1) and we only find a small fraction of the MRB to be susceptible (TI < 0.5) to this compound (Fig. 2C). This, together with the 14x higher amounts of MBOA in the rhizosphere (Table S1) and the main finding that MRB exhibited the broadest range of tolerances to MBOA (Fig. 2B), lead us to conclude that MBOA is the most selective BX compound in the rhizosphere.

### Compound-specific growth inhibition of maize root bacteria in vitro

We also examined bacterial tolerance to non-methoxylated compound analogs of MBOA and AMPO: BOA and APO (Table S1). Given the fact that methoxy groups have been reported to increase the reactivity of benzoxazinoids (67), we hypothesized that the non-methoxylated compounds BOA and APO would exert weaker antimicrobial activity on the MRB than their methoxylated relatives MBOA and AMPO. For BOA, most strains were tolerant (16/52 strains) or moderately tolerant (30/52 strains) while only 6 strains were susceptible (Fig. S7A, Fig. S8A). Hence, bacterial tolerance was generally higher to BOA compared to MBOA (Fig. 2B). Similar taxonomic groups were susceptible (Rhizobiaceae) or tolerant (Bacillaceae) to both BOA (Fisher exact test: tolerance group ∼ family, p < 0.001) and MBOA (Fig. 2B). This finding was further supported by a significant positive correlation of the TIs of all MRB isolates to these two related compounds (Fig. S7B). The spectrum of bacterial tolerance to APO included a large proportion of tolerant MRB strains (37/52 strains), a few moderately affected (7/52 strains), and a few susceptible isolates (8/52; Fig. S7C, Fig. S8B). Overall, bacterial tolerance was lower to APO than AMPO (Fig. 2C), indicating that APO is more toxic than its methoxylated relative AMPO. Again, we find similar taxonomic groups to share APO (Fisher exact test: tolerance group ∼ family: p < 0.05) and MBOA tolerances, which were significantly correlated (Fig. S7D). The positive correlations of bacterial tolerances between the non- and methoxylated compound pairs (BOA/MBOA and APO/AMPO) suggest similar modes of toxicity and tolerance. Bacterial tolerances did not correlate between the non-methoxylated compounds BOA and APO (Fig. S7E). In summary, we found higher bacterial tolerance to non-methoxylated BOA (compared to MBOA) while the MRB tolerated the non-methoxylated APO less than AMPO. Hence, the methoxy group (67) increases the toxicity of (M)BOA but not the aminophenoxazinones A(M)PO.

### Root bacteria from non-BX-exuding plants are less BX-tolerant

Having identified that MRB selectively tolerate the most abundant compound accumulating in the rhizosphere (Fig. 2B, Table S1), we hypothesized that bacteria from a non-BX-exuding plant should be not adapted to BXs and therefore, be less tolerant to BXs than bacteria isolated from a BX-exuding host. To test this hypothesis, we mapped the 16S rRNA sequences from MRB to the strain collection of Arabidopsis bacteria (AtSphere) (57). We selected the most similar bacteria and tested them for tolerance to MBOA. We found that most strains (33/53) were clearly susceptible to MBOA and that 16/53 were moderately affected (Fig. 3A, Fig. S9A). Only 4/53 AtSphere strains were tolerant to MBOA compared to 12/52 tolerant MRB strains (Fig. 2B). Comparing the taxonomic groups of both collections, we found the AtSphere strains of the families Enterobacteriaceae, Microbacteriaceae, Pseudomonadaceae, and Sphingomonadaceae to be less tolerant to MBOA than the corresponding MRB strains (Fig. 3B, Fig. S9B). In the MRB collection, strains of these families showed higher tolerance to MBOA. For the other families, we found no differences in tolerance between the two collections, presumably because these groups are tolerant (Bacillaceae) or susceptible (Micrococcaceae and Rhizobiaceae) to MBOA *per se*. These results show that bacteria isolated from a BX-exuding host are generally more tolerant to MBOA than bacteria isolated from a non-BX-exuding plant, which is suggestive of adaptation to host-specific exudate metabolites.

**Figure 3:**
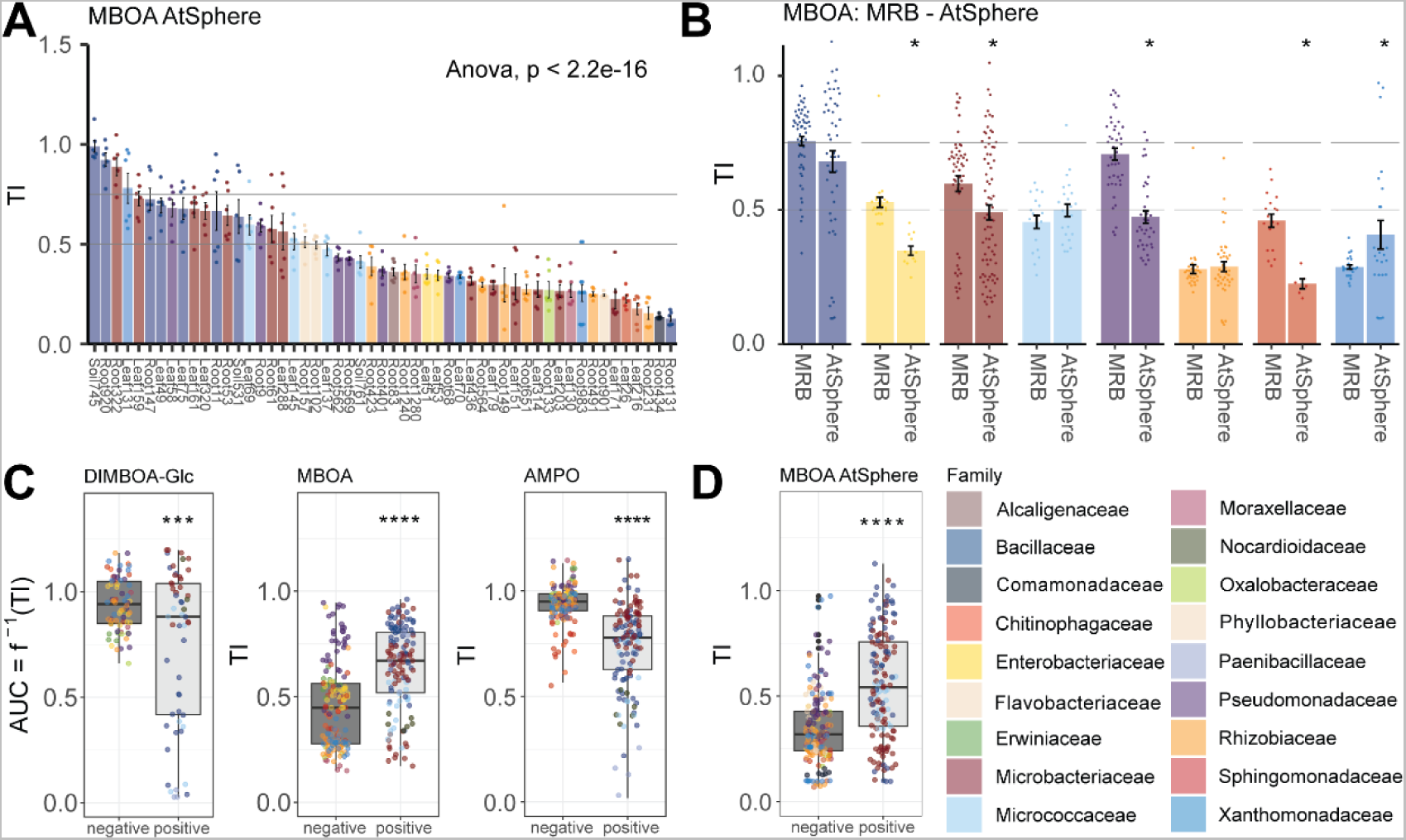
Host-specific tolerance of plant bacteria and cell wall structure defining tolerance. **A)** Isolates of the AtSphere collection are colored according to family taxonomy and their tolerance index (TI) for MBOA is reported. **B)** Tolerance indices of MRB and AtSphere, summarized across families. **C)** Growth and tolerance values summarized for in DIMBOA-Glc (AUC), MBOA (TI), and AMPO (TI) of gram-positive and gram-negative MRB strains and **D)** AtSphere strains. Bar graphs (A and B) show means ± SE bar with individual data points (n = 6 for MBOA). Results of pairwise t-test and ANOVA are shown inside the panels, p-value < 0.05 = *.

### Bacterial tolerance to BXs is related to their cell wall structure

Since we found taxonomically related strains to have similar tolerance levels to different BX compounds (Fig. 3), we tested whether tolerance depended on phylogenetically distinctive features such as cell wall structure. Cell walls of gram-positive bacteria are characterized by a thick peptidoglycan layer while gram-negative bacteria have thin peptidoglycan layers located between an inner and an outer membrane. Gram-positive MRB isolates were significantly more tolerant to MBOA (Fig. 3C) and BOA (Fig. S7A) compared to the gram-negative ones. We found the opposite for AMPO (Fig. 3C) and APO (Fig. S7F), as well as for DIMBOA-Glc (Fig. 3C) with gram-negative bacteria being more tolerant. Consistently, gram-positive AtSphere strains were also more tolerant to MBOA than gram-negatives (Fig. 3D). Together this suggests that cell wall structure can partially explain the tolerance patterns of the different bacteria to BXs and aminophenoxazinones.

### Bacterial tolerance to MBOA explains BX-dependent abundance on maize roots

To test the hypothesis that bacterial tolerance to BXs explains the BX-dependent structuring of maize root microbiomes, we squared our *in vitro* BX tolerance data with microbiome profiles of BX-exuding wild-type and BX-deficient *bx1* maize lines. These microbiome profiles were from the same plants from which most of the MRB strains were isolated from (26). After mapping the 16s rRNA sequences of the MRB strains to the operational taxonomic units (OTUs) of the sequencing data, we first tested for differences in their mean abundances between wild-type and *bx1* roots and rhizospheres (Fig. S10). OTUs of the Bacillaceae and Microbacteriaceae were generally enriched on wild-type plants, while Xanthomonadaceae and Rhizobiaceae OTUs were depleted - a finding, reminiscent of the *in-vitro* tests where these families contained BX-tolerant or non-tolerant strains (Fig. 2). Next, we correlated the tolerance indices of the MRB strains in the different BX compounds with the abundance changes of their corresponding OTUs on wild-type vs. *bx1* microbiomes. Our specific focus lay in determining which BX compounds would best explain the differential abundances of the OTUs. Bacterial tolerance to MBOA correlated best and significantly positively with BX-dependent abundance in the root and rhizosphere microbiomes (Fig. 4A & B). This means that MBOA-tolerant strains mapped to abundant OTUs on BX-exuding roots and their rhizospheres while susceptible strains correspond to low abundant OTUs. Fig. 4C summarizes the comparisons for the other tested compounds. Bacterial tolerance to DIMBOA-Glc does not correlate with BX-dependent abundance (Fig. S11A). A positive correlation was found for bacterial tolerance to BOA which is weaker than MBOA (Fig. S11B). In contrast, bacterial tolerance to AMPO correlated negatively with BX-dependent abundance in the root and rhizosphere microbiome, while tolerance to APO did not correlate (Fig. S11C & D). It is noteworthy that the strongest positive correlation was found for MBOA, which is both the most abundant compound accumulating in the rhizosphere (Table S1) and the compound to which we found the broadest range of bacterial tolerances (Fig. 2B). We conclude that tolerance to MBOA explained, at least partially, the bacterial abundance on BX-exuding vs. BX-deficient roots and that MBOA – with its antibiotic activity – acts as a key driving factor to structure the maize root microbiome.

**Figure 4:**
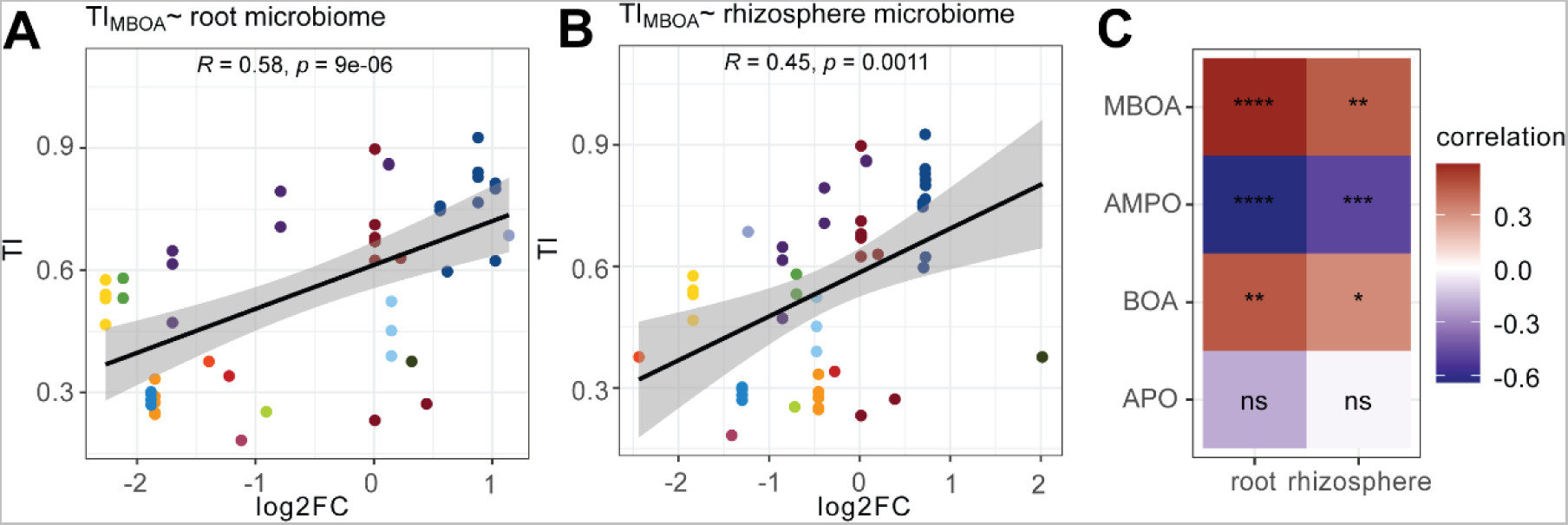
Correlation of bacterial BX tolerance with BX-dependent abundance on maize roots. **A)** Correlation between TIs of all MRB to MBOA with the log2fold changes (log2FC; comparing wild-type vs. bx1 plants) of their corresponding OTUs in root and **B)** in rhizosphere microbiome profiles. Pearson’s product-moment correlation test was performed. R, correlation coefficient; p, p-value. **C)** Heatmap with correlation coefficients R between TIs in MBOA, AMPO, and the non-methoxylated BOA and APO and log2FC of abundances on WT to bx1 roots. Asterisks indicate significance levels, p-value < 0.0001 = ****, < 0.001 = ***, < 0.01 = **, 0.05 = *)

## Discussion

Maize plants shape the composition of their root microbiome through root benzoxazinoid exudation (35–38). To investigate the underlying mechanisms, we established a strain collection of maize root bacteria (MRB) and tested their tolerance to BXs using *in vitro* growth assays. We show that BXs selectively act as antimicrobials on members of the maize root microbiome. Further, we find that bacterial tolerance to the major BX in the rhizosphere (MBOA) correlates with their abundance on BX-exuding maize roots. Below, we discuss the underlying mechanisms and biological implications of our findings.

### Benzoxazinoids act as selective antimicrobials and are better tolerated by bacteria isolated from a BX-exuding host plant

Various plant specialized metabolites have been studied for their selective antimicrobial activities against root microbes. However, such metabolite-microbe interactions are rarely interpreted for native vs. non-host contexts. Non-host refers to context where the root microbes and root metabolites do not originate from the same host. Non-host examples include, that coumarins inhibit the growth of the bacterial pathogen *Ralstonia solanacearum* (isolated from tobacco leaves) (68, 69) or that beneficial *Pseudomonas simiae* (isolated from wheat rhizosphere) and *Pseudomonas capeferrum* (isolated from potato rhizosphere) tolerate high levels of coumarins, compounds that are not produced by tobacco, wheat or potato (31). Also the metabolite-microbe interaction of BXs and Arabidopsis bacteria (59) is a non-host example. Here, the non-methoxylated BXs compounds BOA and APO, both specialized metabolites of wheat and rye, were found to selectively inhibit the growth of Arabidopsis root bacteria in a strain- and compound-specific manner. Fewer studies considered native host context where root metabolites and root microbes belong to the same plant species. A native host example is the study of Arabidopsis root bacteria for their tolerance to coumarins (30). The highly antimicrobial coumarin fraxetin inhibited the growth of bacterial strains belonging to the Burkholderiaceae, while this was not the case for the less toxic scopoletin.

For our study, testing maize and Arabidopsis bacteria for their tolerance against the specialized metabolites of maize, we specifically considered both native and non-host contexts. The native host screening revealed selective growth inhibition of maize root bacteria in an isolate- and compound-dependent manner (Fig. 2). We found highly selective antimicrobial activities of the different BX compounds tested, even with strains that are tolerant to high concentrations of the most abundant and selective BX in the rhizosphere (MBOA). Tolerant strains mostly belonged to the gram-positive Bacillaceae, while gram-negative Xanthomonadaceae and Rhizobiaceae were generally susceptible to MBOA (Fig. 3C). This general pattern of tolerant gram-positive and susceptible gram-negative bacteria was consistent with our non-host screening of Arabidopsis bacteria for MBOA tolerance (Fig. 3D). Also, the general conclusion of selective growth inhibition in an isolate-dependent manner applied to the Arabidopsis bacteria. Importantly, the direct comparison by host context revealed that the ’non-host’ bacteria from Arabidopsis were generally less tolerant to the specialized compounds of maize than their ‘native host’ counterparts isolated from maize (Fig. 3B). This finding suggests that members of the root microbiome have adapted to the antimicrobial root exudates of their host plant. Further research is required to test if our results can be generalized, for instance by screening the AtSphere (60) and MRB (this study) collections for their tolerance against specialized Arabidopsis compounds like coumarins. If adaptation to host antimicrobials were broadly applicable, this would imply that host-specific mechanisms have evolved in the genomes that enable root bacteria to tolerate host-specific substances. Therefore, genomic studies, e.g. comparing the genomes of Arabidopsis and MRB, should allow the identification of the underlying genetic components for tolerance to host-specific antimicrobial compounds.

### Cell wall structure defines BX tolerance

We found, both in ‘native host’ and ‘non-host’ contexts, that gram-positive bacteria were more tolerant to MBOA than gram-negative bacteria (Fig. 3CD), indicating that cell wall properties are important for bacterial tolerance to BXs. For the tolerant gram-positive bacteria, this suggests that the thick peptidoglycan layer of the cell wall presents a first level of protection, possibly by preventing the entry of MBOA into the cell. Seemingly consistent, the cell wall of gram-negative bacteria consists of a thin peptidoglycan layer. This thin peptidoglycan layer is located between an inner and an outer membrane and possibly the exposure of the membranes explains the susceptibility of gram-negative bacteria to BXs. Little is known about the mode of action of BXs against microorganisms (67, 70). Suggested mechanisms of BX toxicity include intercalation with DNA, chelation, or active import into microbial cells due to their siderophore-like function. For the overall activity, the lipophilicity of BXs, which influences diffusion across cellular membranes, is important. Our work supports this with the finding of the opposed tolerances of bacteria to the less lipophilic (M)BOA versus the more lipophilic A(M)PO (Fig. 3C & S7E). Hence, we infer that bacterial membranes are key factors for BX toxicity.

### Mechanisms of BX tolerance

There is a key mechanistic question emerging from the finding of opposed tolerances of gram-negative vs. gram-positive bacteria: How do generally susceptible gram-negative bacteria gain tolerance to BXs? Options for increased tolerance are related to the transport of the compounds across the membranes, either by blocking their import (if an active process) or by exporting them out of the cell. The recent study by Machado et al. (2020) provided the first mechanistic insights from studying how the gram-negative *Photorhabdus* bacterium tolerates MBOA (58). Analysis of isolates with increased MBOA tolerance which were selected after experimental evolution in the presence of MBOA revealed multiple membrane-related mechanisms for BX tolerance. For instance, a mutation in the aquaporin-like membrane channel aqpZ conferred MBOA tolerance, which suggests a mechanism related to preventing BX import. Alternative to transport, a third possible mechanism for increased tolerance would be to metabolize the compounds. More in-depth studies are needed to uncover the underlying mechanisms of bacterial tolerance to BXs. Again, strain collections like the AtSphere (60) and MRB (this study), will be powerful resources to study e.g. the BX metabolizing traits root bacteria using phenotypic and genomic strain comparisons.

### Tolerance to specialized compounds explains community structure

Root microbiome structure is thought to arise from plant and microbial processes, both involving small metabolites. Microbes, for instance, contribute to community structure by competing with their microbiome peers based on specialized exometabolites. Such antimicrobial compounds were recently shown to act as competence determinants for *Pseudomonas* bacteria in the root microbiome (27). Bulgarelli et al. (2013) proposed a two-step selection model for the plant processes with the central idea that root exudates ’fuel an initial substrate-driven community shift in the rhizosphere, which converges with host genotype–dependent fine-tuning’ of the root microbiome (1). The first step is relatively well understood, where the microbial recruitment from the surrounding soil, also known as the ‘rhizosphere effect’, is mostly fueled by primary metabolites that function as carbon substrates for microbial growth. For step two, plant specialized metabolites were repeatedly identified as key factors for microbiome assembly among several other drivers (15, 16). Typically, these conclusions stem from shifts in microbiome profiles of mutants that lack certain specialized metabolites compared to wild-type plants (22, 31, 32, 35–38). However, the mechanisms by which plant specialized metabolites shape microbiome composition and the processes that regulate microbial community structure remain poorly understood. Here we provide evidence that bacterial tolerance to plant specialized metabolites explains how these compounds shape the root microbiome. We systematically determined the tolerances of MRB to BXs (Fig. 2 & S7) and found that bacterial tolerance to MBOA, the most abundant antimicrobial metabolite in exudates of their host plant, correlated significantly with their abundance on benzoxazinoid-exuding roots (Fig. 4). Thus, our results indicate that MBOA- tolerance is an important trait for abundant colonization of maize roots, even if further experiments would be required to validate this finding. This could include the use of synthetic communities composed of strains with different benzoxazinoid tolerance levels and exposing them to different BXs or inoculate axenic roots of wild-type and mutant maize with BX tolerant and susceptible strains and measure the benzoxazinoid-dependent colonization.

### Linking bacterial tolerance to the establishment of a healthy root microbiome

Many specialized metabolites of plants have dual functions, both suppressing pathogens and recruiting beneficials. Thus, their exudation is an important tool for plants to steer the establishment of a healthy root microbiome (71). Such dual functions are also known for coumarins and BXs. Scopoletin, the dominant coumarin in the rhizosphere of Arabidopsis, inhibits the soil-borne fungal pathogens *Fusarium oxysporum* and *Verticillium dahlia* while promoting *P. simiae* and *P. capeferrum*, both beneficial rhizobacteria (31). Similarly, BXs inhibit fungal pathogens *Setosphaeria turtica*, *Exserohilum turcicum,* and *Fusarium spp.* (43) and reduce the virulence of the phytopathogenic bacterium *Agrobacterium tumefaciens* (55). At the same time, BXs enrich beneficial bacteria (14, 31, 54, 71, 72) by acting as chemoattractant for the beneficial rhizobacterium *Pseudomonas putida* to maize roots (54). Although most of these examples are from ‘non-host’ contexts, they suggest that plants assemble a health-promoting root microbiome with the exudation of their specialized compounds. The established MRB collection allows us to study this also in a native host context. Here, the screening of the MRB strains for BX tolerance revealed the first indications towards the establishment of a healthy root microbiome with ‘pathogen suppression’ and ‘recruitment of beneficials’. Suppression of pathogen implies their susceptibility to BXs, which is what we have observed for all *Agrobacterium* strains in the collection (Fig. 2). Analogously, recruitment of beneficial implies that they tolerate the BXs, which is what we see for isolates of *Pseudomonas* and *Bacillus* (Fig. 2), both families with known beneficial plant bacteria. Now, functional work e.g., exploring the functional plant phenotypes of the MRB strains, will corroborate the links between susceptibility and tolerance to BXs with ‘pathogen suppression’ and ‘recruitment of beneficials’, respectively.

In conclusion, based on our results and the general facts that many specialized metabolites of plants have an antimicrobial function (22, 30–32, 67) and that many of them are key factors shaping community structure (22, 31, 32, 35–38), we propose that bacterial tolerance to plant-secreted antimicrobials is a mechanism that determines host-specific microbial community composition.

## Materials and Methods

### Quantification of root bacterial community size

We quantified the size of the bacterial community on roots of B73 and *bx1* mutant plants from two independent experiments (Table S2) by plating the cultivable bacteria and by quantitative PCR. See the Supplementary Methods and Table S2 for details.

### Establishment of MRB culture collection

The culture collection of maize root bacteria (MRB, Dataset S1) was built with strains isolated in five independent experiments (Table S2). All strains were isolated from greenhouse pot experiments with Changins soil, i.e., batches of the same soil where we first demonstrated the microbiome structuring activity of BXs (37). Most strains originate from wild-type maize plants (inbreed line B73) and a small subset of strains was isolated from BX-deficient *bx1*(B73) plants. Most strains were isolated from ‘dirty roots’ including the root and the rhizosphere fraction (marked as ‘RoRh’, Dataset S1), and a few strains were isolated from washed roots, rhizosphere, or soil extracts (described in Supplementary Methods). The extracts were diluted from 1:10^-3^ to 1:10^-6^ in 10 mM MgCl_2_ for plating so that spreading of 50 μl extract with a delta cell spreader (Sigma-Aldrich, St. Louis, USA) resulted in a density of 100-300 CFU on square plates (12 x 12 cm, Greiner bio-one, Kremsmünster, Austria). The media used for isolation are listed in Table S4. The plates were incubated at room temperature (22 - 24 °C) for 5-10 days, single colonies were picked and re-streaked on full-strength TSB or LB media until the isolates were visibly pure colonies. Bacterial strains were routinely sub-cultured at 25 °C – 28 °C in tryptic soy broth (TSB, Sigma-Aldrich) or Luria-Bertani medium (LB, Carl Roth, Karlsruhe, Germany) liquid or solid medium amended with 15 g/l agar (Sigma-Aldrich).

For cryopreservation, single colonies of pure strains were inoculated in full-strength liquid TSB or LB medium, grown for two days at 28°C with 180 rpm shaking, and then mixed with the same volume of 40% sterile glycerol (Sigma-Aldrich) in single screw cap microtubes (Sarstedt, Nürnbrecht, Germany). The resulting 20% glycerol stocks were slowly frozen down and stored at −80°C. The same liquid cultures, of which the glycerol stocks were prepared, were used for Sanger sequencing-based isolate identification using the 16S rRNA gene as described in Supplementary Methods and Table S3. All sequences together with taxonomies and metadata of the MRB are listed in Dataset S1 and the strain collection is maintained in the laboratory of the authors.

We identified the MRB strains among operational taxonomic units (OTUs) or amplicon sequence variants (ASVs) of published 16S rRNA gene profiles of maize root communities (35–37). See the Supplementary Methods for a detailed description of the mapping method. The sequence similarities of the strains to the microbiome members (ASVs) are listed in Dataset S2.

Phylogenetic analysis was performed with the Sanger-based 16S rRNA gene sequences. They were first concatenated, then aligned using MAFFT v. 7.475 (73) with default options, and analyzed with RAxML v. 8.2.12 (74). The multi-threaded version ‘raxmlHPC-PTHREADS’ was used with the options ‘ -f a -p 12345 -x 12345 -T 23 -m GTRCAT’ with 1000 bootstrap replicates. The phylogenetic tree was visualized (Fig. 1) and annotated in R (package ggtree, 102).

### High-throughput growth phenotyping of MRB strains

To screen MRB strains for their tolerance against various BXs and degradation compounds we developed a custom, high-throughput, in vitro, liquid culture based-growth system (76). In brief, many bacteria strains are cultured in parallel and in a replicated manner in many 96-well plates, which are handled with a stacker (BioStack 4, Agilent Technologies, Santa Clara, United States), so that the connected plate reader (Synergy H1, Agilent Technologies) records bacterial growth via optical density (OD600, absorbance at 600 nm) over time. In 50% TSB (Table S4), we tested the following compounds and concentrations: DIMBOA-Glc (2500 µM), purified from maize seedlings (purification method is detailed in the Supplementary Methods), synthetic MBOA and BOA (at 250, 500, 625, 1’250, 2’500 and 5’000 μM), AMPO (10, 25 and 50 μM), synthesized in our lab following a published procedure (77) and synthetic APO (10, 25, 50 and 100 μM, Table S5). We included controls with the solvent DMSO (Sigma-Aldrich), of which the concentration was kept constant in all treatments. We set up separate runs for the different compounds and in one run, we always tested all concentrations of a compound against 52 MRB strains. Every compound was repeated in at least 2 runs. The assay setup is further detailed in the Supplementary Methods. The bacterial growth data were analyzed in R (version 4.0, R core Team, 2016). For growth we calculated the area under the curve (AUC; function *auc()* from package MESS, (78) and normalized growth in a treatment relative to the control. Such normalized bacterial growth data of a given concentration was statistically assessed (compound vs control) using one-sample t-tests (p-values adjusted for multiple hypothesis testing). As a measure of tolerance of a given strain to a given compound, a specific tolerance index (TI) was calculated from the normalized AUC values across all tested concentrations of that compound. This calculation uses again the *AUC ()* function taking the AUC across the normalized AUC values across all tested concentrations. In contrast to defining medium inhibitory concentrations (IC50), the TI method is more broadly applicable as it allows tolerance comparisons including strains that are not inhibited at the highest tested concentration. The TI ranges from 1 (full tolerance, no growth inhibition at highest tested concentration) to 0 (full susceptible, no growth at lowest tested concentration) and we classified strains as tolerant (TI ≥ 0.75), intermediate (0.75 > TI ≥ 0.5) or susceptible (TI < 0.5). TI variation was assessed across strains using analysis of variance (ANOVA, TI ∼ Strain) or across tolerance classes testing for a taxonomic signal using a Fisher’s exact test (TI class ∼ family). TIs were compared between different compounds using Pearson correlation. To test whether bacterial tolerance to BXs explains the BX-dependent structuring of maize root microbiomes (Fig. 3), we analyzed the TI data relative to the microbiome data of wild-type and *bx1* mutant maize (37). After mapping the MRB strains to the sequencing data (Supplementary Methods), we determined differential colonization of the corresponding OTU on wild-type vs. *bx1* roots and rhizospheres. Next, we correlated strain TI with differential colonization (log_2_FC) of corresponding OTUs using Pearson and its product-moment test. The code used for statistical analysis and graphing is available from https://github.com/PMI-Basel/Thoenen_et_al_BX_tolerance. The following further R packages were used: Tidyverse (79), Broom (80), DECIPHER (81), DESeq2 (82), emmeans (83), ggthemes (84), multcomp (85), phyloseq (86), phytools (87), vegan (88) in combination with some custom functions.

### Bacterial genomes

We generated the genomes of the MRB strains in four efforts (Dataset S1). The genomes were generated using PacBio and Illumina as detailed in the Supplementary Methods. To assemble the genomes, we used similar pipelines as described in detail in the Supplementary methods. The raw sequencing data and the genome assemblies and annotations have been deposited in the European Nucleotide Archive (http://www.ebi.ac.uk/ena) with the study accession XYZ (sample IDs ABC to DEF) (Dataset S1).

## Acknowledgments

We thank Prof. Dr. Julia Vorholt (ETH Zurich) and Prof. Dr. Paul Schulze-Lefert (MPMI Cologne) for sharing the bacterial strains from the AtSphere collection. Big thank goes to Niklas Schandry (GMI Vienna, later LMU Munich) for the introduction of the high-throughput chemical phenotyping method and the sharing of the code for the analysis of the bacterial growth curves. Further we thank Dr. Pamela Nicholson (next-generation sequencing platform University of Bern) for technical support with sequencing and Kerstin Schneeberger (Agroscope Zurich) for the help with the initial genome assembly of Set 1. We thank Selma Cadot for the preparation of the root DNA samples used for qPCR, Corinne Suter for the technical assistance with DNA extraction and culturing of bacteria, Sandro Rechsteiner for the isolation of the initial maize root bacteria strain collection, and Florian Enz for the plant maintenance. This work was supported by the Interfaculty Research Collaboration “One Health” of the University of Bern (www.onehealth.unibe.ch). This work was also supported by MINECO, Spain, RYC-2015–19154 and grant SEV-2015-0533 funded by MCIN/AEI/10.13039/501100011033, and by the CERCA Programme / Generalitat de Catalunya to I-R-S.

## Supporting Information

### Supplementary Methods

#### Bacterial extracts

Various types of bacterial extracts were prepared for establishing the culture collection and quantifying root bacterial community size. First, bacterial extracts from ‘dirty roots’ (marked as ‘RoRh’ in Dataset S1) were prepared from 10 cm long root fragments (corresponding to the depth of −1 to −11 cm in soil) that were chopped into small pieces with a sterile scalpel after shaking off loose soil. These root fragments with firmly attached rhizosphere soil were then placed into 50 mL centrifuge tubes containing 10 mL sterile magnesium chloride buffer and Tween20 (10 mM MgCl_2_ + 0.05 % Tween; Sigma-Aldrich) for homogenization with a laboratory blender (Polytron, Kinematica, Luzern, Switzerland; 1 minute at 20’000 rpm) followed by additional vortexing for 15 seconds. Extracts of washed roots (marked as ‘root’ in Dataset S1) were prepared analogously, except that the roots were washed twice in 50 mL centrifuge tubes with 25 mL of sterile deionized water and shaking the tubes 30 times vigorously to wash off the rhizosphere before cutting them in small pieces for homogenization. The rhizosphere fraction of the washing step was pelleted by sedimentation (supernatant was discarded) and resuspended in 5 mL MgCl_2_-Tween (10 mM, 0.05%) to prepare the rhizosphere extracts for plating (marked as ‘rhizo’ in Dataset S1). Plating extracts from soil (marked as ‘soil’ in Dataset S1) were prepared by mixing 5 g of soil from the pot experiment with 5 mL MgCl_2_-Tween (10 mM, 0.05%) and vortexing for 15 seconds.

#### Quantification of root bacterial community size with plating

We quantified the sizes of the root bacterial communities of B73 and *bx1*(B73) (1) plants in two greenhouse experiments (Table S2). In both experiments, one half of the roots was freshly used for plating the cultivable bacteria and the other half stored at −80 °C for culture-independent qPCR analysis (see below). The first experiment consisted of 6-week-old plants and 7-week-old plants (same as ‘Isolation 4’) were analyzed in the second experiment. Extracts of washed roots were freshly prepared as described above, serially diluted for plating and 20 µL were plated on 10 % TSA (tryptic soy medium amended with 15 g/L agar; both Sigma-Aldrich) plates containing cycloheximide (10 mg/L, Sigma-Aldrich). Plates were tilted to spread the 20 µL drops for counting, then incubated for six days at room temperature. The forming colonies were counted, multiplied by the dilution factor and the volume plated, and then normalized with the sample fresh weight. The colony forming unit (CFU) data was transformed with log10 prior to statistical analysis (T-test) and visualization.

#### Quantification of root bacterial community size with qPCR

Complementary to CFU plating, we quantified bacterial community size on the second half of root samples using qPCR analysis. The frozen roots were lyophilized, and DNA was extracted using the Nucleo-Spin Soil DNA extraction kit (Macherey-Nagel, Düren, Germany) following the manufacturer’s protocol. Additionally, we also utilized available DNA samples of our previous field experiments in Changins (CH, (2)), Reckenholz (also CH) and Aurora (US, both (3)) for qPCR analysis. For all DNA samples, the concentration was measured using the AccuClear® Ultra High Sensitivity dsDNA Quantification Kit (Biotium, Fremont, United States) and adjusted to 1 ng/μL. qPCR reactions were set up in a total volume of 20 μL containing HOT FIREPol EvaGreen qPCR Mix Plus (Solis Biodyne, Tartu, Estonia), 250 nM of each primer, 0.3% bovine serine albumin, and 10 ng of root DNA. The size of the bacterial community was quantified on genomic DNA based on the bacterial 16S rRNA gene (primers 799F and 904R, Table S3) relative to the maize gene Actin (primers ZmActin1_F and ZmActin1_R1, Table S3). No template control reactions containing water were run in parallel as negative controls. qPCR reactions were set up (in triplicates for greenhouse experiments and in single reactions for samples from field experiment) using the Myra Liquid Handler (Bio Molecular Systems, Upper Coomera, Australia) and ran on a CFX96 Real Time System (Bio Rad, Hercules, California). The cycling program included an initial denaturation step at 95 °C for 15 min, followed by 80 cycles of 95 °C for 15 s, 63 °C for 40 s and 72 °C for 20 s, a hold phase at 72 °C for 10 min, followed by melting curve analysis (temperature incrementally increased by 0.5°C from 65 to 95 °C with steps held for 5 s). Raw data were exported directly from Bio-Rad CFX Manager 3.1 and imported into LinRegPCR version 2016.0 (4) to determine cycle threshold (Ct) and efficiency (E) using the default baseline limit option. The bacterial 16S rRNA gene signal was normalized to the plant signal using the following formula: 16S rRNA/plant gene = E_plant gene_^Ct_plant gene_/E_16S_^Ct_16S_, where Ct values of the individual reactions and mean E values over all reactions of a given primer pair and run were used for calculation (5). Data was transformed with log2 prior to statistical analysis (T-test) and visualization.

#### MRB isolate identification

The taxonomy of the purified isolates of the maize root bacteria (MRB) collection were identified by sequencing parts (base pairs 27 to 1492) of the 16S rRNA gene using Sanger technology. Liquid cultures were diluted 1:10 or 1:100 in sterile water and used as template for PCR. The PCR reactions were set up as follows: 15 μL sterile water, 15 μL 2x DreamTaq buffer (Thermo Fisher Scientific, Waltham, USA), 1.5 μL of each primer (stock concentration 10 μM, 27f and 1492r; sequences in Table S3) and 2 μL of the diluted liquid culture as DNA template. For some bacteria, the DNA was extracted using the GenElute™ Bacterial Genomic DNA Kit (Sigma-Aldrich). PCR was performed in a Biometra T-advanced cycler according to the following program: 95°C for 3 min, 30 cycles with 95°C for 15 s, 55°C for 15 s and 72°C for 45 s followed by final elongation at 72°C for 5 min. PCR products were verified on an agarose gel (1 %; Sigma-Aldrich) and sent for Sanger sequencing with the primers 1492r and/or 27f (Microsynth, Balgach, Switzerland). Sanger sequences were blasted against the NCBI database (National Center for Biotechnology Information, Rockville Pike, USA) for species identification. All metadata, sequences, and taxonomies of the MRB culture collection are listed in the Dataset S1 and the strain collection maintained in the laboratory of the authors.

#### Mapping MRB isolates to microbiome profiles

##### Rationales

We mapped the 16S rRNA gene (Sanger) sequences of the MRB strains to the 16S rRNA gene sequences of published maize community profiles (i.e., to the OTUs, ZOTUs or ASVs of these datasets). The first purpose was to investigate abundance of community members corresponding to MRB strains in profiles of maize roots from where the strains were isolated from. For this we mapped the MRB sequences to the data of the feedback experiment reported in Hu et al. 2018 (2). This was a greenhouse experiment with pots filled with natural field soil from the Changins site. Second, to study presence and abundance of community members (corresponding to MRB strains) in root profiles of field grown maize, we mapped the MRB sequences to the datasets of Changins (the field experiment data reported), Reckenholz and Aurora (3) and the pot experiment in Sheffield (6). The third purpose was to examine differential abundance of the MRB strain corresponding community members in profiles of BX-producing vs BX-deficient plants (the field data of Changins (2) also includes profiles of the *bx1* mutant maize line).

##### Bioinformatics

Because the published datasets (2, 3, 6) utilized different bioinformatic approaches, we re-processed the deposited raw sequence data to have uniformly analyzed microbiome data, to which we then mapped the MRB strains (see below). The raw sequence reads were quality checked with FastQC (7), demultiplexed by cutadapt (8) and then processed using the DADA2 pipeline with default options (9). The sequences were filtered by allowing maximal expected errors of two and with maximal zero Ns. Reads were truncated at the first instance of a quality score of less than three. The forward reads of the Changins, Reckenholz and Aurora data were trimmed to 250 bp and reverse reads to 170 bp. As the sequences of the Sheffield data were only 250 bp long, forward reads were not trimmed and reverse reads to 200 bp. Shorter reads were discarded. For each MiSeq run, a parametric error model was learned by the DADA algorithm and inferred to the previously dereplicated samples. Then the forward and reverse reads were merged if the overlap was identical and at least twelve bases long. A single (ASV) table was created as all datasets used the same 16S rRNA gene primers. We removed chimeras and assigned taxonomy to the amplicon sequence variants (ASVs) with the naive Bayesian classifier method (9) and the SILVA database (10). Scripts are available from https://github.com/PMI-Basel/Thoenen_et_al_BX_tolerance. The computations were performed at the Vital-IT (https://www.vital-it.ch) center for high-performance computing of the SIB Swiss Institute of Bioinformatics and at sciCORE (http://scicore.unibas.ch/) scientific computing center at University of Basel.

##### Mapping

We aligned the 16s rRNA sequences of the MRB obtained by Sanger sequences to overlap with the 16S rRNA gene region (primers 799F and 1193R; Table S3) of the microbiota profiles using the function *AlignSeqs* (R package DECIPHER, 99). Then, a distance matrix was calculated for all MRB sequences to the identified amplicon sequence variants (ASVs) of the respective datasets using the function *DistanceMatrix* (DECIPHER). We did not consider mappings with <97% sequence similarity, the typical threshold for defining operational taxonomic units (OTUs). Strains typically mapped to several ASVs (within 97%), hence their summed relative abundance was taken. The similarities of the MRB strain sequence to the microbiome members (i.e., the ASVs) are listed in Dataset S2. Scripts are available from https://github.com/PMI-Basel/Thoenen_et_al_BX_tolerance. Calculations were performed at sciCORE (http://scicore.unibas.ch/) scientific computing center at University of Basel.

#### Isolation and purification of benzoxazinoids form maize plants

DIMBOA-Glc was isolated from maize plants as described below. Ca. 200 g of maize leaves (*Zea mays*, Akku) were frozen and ground in liquid nitrogen. The resulting powder was placed in 1.5 L MeOH and allowed to warm up to room temperature. The resulting suspension was homogenized with an immersion disperser (PT-1035, Kinematica AG, Malters, Switzerland) and filtered through a P3 sintered glass filter equipped with two layers of filter paper, with suction. The filter cake was collected and suspended again in 0.6 L MeOH. After a second homogenization and a new filtration, the filtrates were combined and concentrated under reduced pressure with a rotary evaporator (RC900, KNF Neuberger AG, Balterswil, Switzerland). The aqueous residue obtained was lyophilized with a freeze-drier (LyoQuest -55, Telstar, Terrassa, Spain) to give 8.39 g of crude dry extract. Five runs of purification with ca. 1.7 g of raw material each were performed on an automated flash column chromatography apparatus (CombiFlash Rf+, Teledyne ISCO Inc., Lincoln NE, USA). Solid loading and 120 g silica cartridges were used. The elution gradient was as follow: 0-13% B over 7 min, 13-16% B over 9 min, 16-35% B over 9 min where A = CHCl_3_ and B = MeOH. The fractions eluting between 19 and 25 min were collected, combined, concentrated under reduced pressure and submitted to new runs of purification. Batches of approx. 250 mg were purified separately with solid loading, 40 g silica gold cartridges, and eluting with 0-15% B over 1.2 min, 15-19% B over 7.2 min, 19-30% B over 1.2 min, 30% B over 3.6 min. The fractions eluting between 11 and 14 min were collected, combined, and concentrated under reduced pressure with a rotary evaporator (RC900, KNF Neuberger AG, Balterswil, Switzerland) to obtain 180 mg of a light-yellow foam (hygroscopic) containing ca. 70% DIMBOA-Glc, 15% DIM_2_BOA-Glc, 15% HMBOA-Glc. The analytical data were in accordance with previous literature: UPLC m/z 194.04 [M−Glc−MeOH]_+_, Mass window: 0.02 Da. Retention time: 1.62 min. HRMS calculated for C_15_H_18_NO_10_ [M−H]^−^: 372.0936, found: 372.0944; ^1^H NMR (300 MHz,CD_3_OD) *δ* 7.26 (d, *J* = 8.8 Hz, 1H), 6.75 (d, *J* = 2.6 Hz, 1H), 6.70 (dd, *J* = 8.8, 2.7 Hz, 1H), 5.91 (s, 1H), 4.67 (d, *J* = 7.8 Hz, 1H), 3.78 (s, 3H), 3.9-3.1 (m, 6H) (12).

#### High-throughput growth phenotyping of MRB strains

We have described our high-throughput chemical phenotyping system, which we have used to screen MRB strains for their tolerance against various BXs compounds, in detail (13). Here we document the specific settings used in this study.

Setting up an assay requires the preparation of liquid pre-cultures in a 96-well format from fresh bacterial cultures on solid media plates. Pre-cultures were prepared by transferring isolate colonies with inoculation needles (Greiner bio-one, Kremsmünster, Austria) to 1 mL of liquid 50% TSB (Table S5) in 2 ml 96-well deep-well plates (Semadeni, Ostermundigen, Switzerland). These pre-culture growth plates were covered with a Breathe-Easy membrane (Diversified Biotech, Dedham, USA) and grown until stationary phase for 4 days at 28°C and 180 rpm.

Assays were set up by inoculating 4 µL of the pre-cultures to 200 µL fresh liquid 50% TSB (Table S5) in 96-well microtiter plates (Corning, Corning, USA) containing the compounds and concentrations to be tested: DIMBOA-Glc (2500 µM), MBOA and BOA (250, 500, 625, 1’250, 2’500 and 5’000 μM), AMPO (10, 25 and 50 μM) or APO (10, 25, 50 and 100 μM). These treatments were prepared by mixing their stock solutions into liquid 50% TSB. Stock solutions were prepared in the solvent DMSO (Sigma-Aldrich) depending on the solubility of the compounds (Table S5) and the DMSO concentration was kept constant in each treatment including the control.

All reactions and replicated plates were pipetted using a liquid handling system (Mettler Toledo, Liquidator 96™, Columbus, USA). All plates had lids and were piled up and inserted to a stacker (BioStack 4, Agilent Technologies, Santa Clara, United States), which was connected to a plate reader (Synergy H1, Agilent Technologies, Santa Clara, United States). Using this system, the optical density (OD600, absorbance at 600 nm) of every culture was recorded every 100 min over 68 hours. Prior to each measurement, the plates were shaken for 120 s. In each plate, wells with 50% TSB were included as ‘no bacteria controls’ and in each run one plate containing only media was included to monitor potential contaminations.

We set up separate runs for the different compounds. In one run, we always tested all concentrations of a compound against all 52 strains with 3 replicates per strain and an empty media control plate. For example, a typical run consisted of a total 23 plates that cover 11 treatments (e.g., 6 concentrations of MBOA + 3 concentrations of AMPO + 2 control treatments; 1 plate per treatment) * 162 cultures (e.g., 52 strains and 2 no bacteria controls, all with 3 replicates; distributed on 2 plates) plus 1 media plate without bacteria. Such a run yielded 1782 single growth reactions. We have performed at least 2 runs for every compound (except DIMBOA-Glc 2500 µM due to low availability of the compound). Data were exported from the software of the plate reader (Gen 5, Agilent Technologies, Santa Clara, United States) and imported into R for data analysis (see main methods).

#### Bacterial genomes

We generated the genomes of the MRB strains in four sets (Dataset S1).

##### Set 1

The first set of MRB strains consisted of the following four bacteria: *Pseudomonas* LPB4.O, *Pseudomonas* LPD2, *Rhizobium* LRC7.O and *Rhizobium* LRH8 (Dataset S1). Genomic DNA was extracted from overnight cultures grown in liquid LB medium (Table S5) using the GeneElute Bacterial DNA kit (Sigma-Aldrich). 10 kb insert libraries were prepared from the genomic DNA (BluePippin size selection) and sequenced on a PacBio (Pacific Biosciences, Menlo Park, USA) RSII instrument (one RSII SMRT cell per strain; P6-C4 chemistry) at the Functional Genomics Centre Zurich (http://www.fgcz.ch). European Nucleotide Archive (http://www.ebi.ac.uk/ena) with the study accession XYZ (sample IDs ABC to DEF).

##### Set 2

The second set of MRB strains consisted of 10 bacteria (Dataset S1). Genomic DNA was extracted as for the first set and used for library preparation using NEBNext® DNA Library Prep Kit (New England Biolabs, Ipswich, USA) following the manufacturer’s recommendations. The libraries were sequenced on a NovaSeq 6000 instrument (paired-end 150 bp reads; Illumina, San Diego, USA) by Sequentia (www.sequentiabiotech.com) together with other samples of that company and the target to produce >1 Gb of data for each library. European Nucleotide Archive (http://www.ebi.ac.uk/ena) with the study accession XYZ (sample IDs ABC to DEF).

##### Set 3

The majority of MRB strains (29 strains; Dataset S1) were sequenced in the third set. Total DNA was extracted using the DNeasy UltraClean Microbial Kit (Qiagen, Hilden, Germany) according to the protocol provided. Quantity, purity, and length of the total genomic DNA was assessed using a Qubit 4.0 fluorometer with the Qubit dsDNA HS Assay Kit (Thermo Fisher Scientific), a DS-11 FX spectrophotometer (DeNovix, Wilmington, USA) and a FEMTO Pulse System with a Genomic DNA 165 kb Kit (Agilent, Basel, Switzerland), respectively. Sequencing libraries were made using an Illumina DNA Prep Library Kit (Illumina, San Diego, USA) in combination with IDT for Illumina DNA/RNA UD Indexes Set C and Tagmentation according to the Illumina DNA Prep Reference Guide. The input DNA was set at 200 ng and 5 PCR cycles were employed to amplify the fragmented DNA. Pooled DNA libraries were sequenced paired end on a NovaSeq 6000 SP Reagent Kit v1 (300 cycles) on an Illumina NovaSeq 6000 instrument. The quality of the sequencing run was assessed using Illumina Sequencing Analysis Viewer (version 2.4.7) and all base call files were demultiplexed and converted into FASTQ files using Illumina bcl2fastq conversion software v2.20. All steps from gDNA extraction to sequencing data generation were performed at the Next Generation Sequencing Platform, University of Bern, Switzerland. The raw sequencing data is available from the European Nucleotide Archive (http://www.ebi.ac.uk/ena) with the study accession XYZ (sample IDs ABC to DEF).

##### Set 4

Several Microbacteria strains (13 strains; Dataset S1) were subjected to PacBio sequencing. DNA was extracted following the GES method (14) from fresh agar plate cultures to ensure good quality DNA with low fragmentation. Briefly, 2-4 mL of each bacterial strain was grown overnight in liquid TSB (Table S3) at 28 °C, centrifuged for 10 min at 12’000 rpm at RT, the media was discarded, and the bacterial pellet was re-suspended in 200 µL TE buffer (10 mM Tris-HCl, 1 mM EDTA, pH 8.0). For cell lysis 500 µL of GES solution (guanidium thiocyanate) was added to each bacterial suspension and incubated for 10 min at RT, before the addition of 250 µL of 7.5 M ammonium acetate. The mixture was gently mixed and incubated on ice for 10 min. Thereafter, 500 µL phenol chloroform isoamyl alcohol mixture, 25:24:1 (Sigma-Aldrich) was added, vigorously mixed, and centrifuged for 15 min at 12’000 rpm at 4 °C. The upper aqueous layer was transferred to a fresh tube and 500 µL of chloroform isoamyl alcohol mixture 24:1 (Sigma-Aldrich) was added, vigorously mixed, and centrifuged for 15 min at 12’000 rpm at 4 °C. Once again, the upper layer of fluid was transferred to a new tube and mixed with 0.7 vol. 100 % isopropanol, mixed well, and stored at -20 °C overnight. Precipitated DNA was recovered by centrifugation at 12’000 rpm for 15 min at 4 °C. The DNA pellet was washed once with 80 % ethanol and twice with 70 % ethanol. The pellet was dissolved slowly in 80 µL water with the aid of heating at 55 °C for 1 h. Prior to SMRTbell library preparation, bacterial genomic DNA was assessed for quantity, quality and purity using a Qubit 4.0 flurometer (Qubit dsDNA HS Assay kit; Thermo Fisher Scientific), an Advanced Analytical FEMTO Pulse instrument (Genomic DNA 165 kb Kit; Agilent) and a Denovix DS-11 UV-Vis spectrophotometer, respectively. Multiplexed SMRTbell libraries were prepared for sequencing on the Sequel exactly according to the PacBio guideline entitled: “Procedure & Checklist – Preparing Multiplexed Microbial Libraries Using SMRTbell® Express Template Prep Kit 2.0” - Part Number 101-696-100 Version 08 (November 2021). Concisely, 1 μg of gDNA in 100 µL was used to shear the gDNA using a Covaris g-TUBE (Covaris, Wolburn, US). Subsequently, the sheared gDNA was concentrated and cleaned using AMPure PB beads. The samples were then quantified and qualified to be in the range of 12- 15 Kb using a Qubit 4.0 flurometer (Qubit dsDNA HS Assay kit, Thermo Fisher Scientific) and an Advanced Analytical FEMTO Pulse instrument (Genomic DNA 165 kb Kit, Agilent), respectively. The rest of the procedure as referenced above was followed including removal of single strand overhangs, DNA damage repair, end-repair & A-tailing, ligation of barcoded overhang adapters and then purification of the library using AMPure PB beads. The libraries were quality controlled using the steps described above and then were pooled using the PacBio microbial multiplexing calculator. Prior to and after size selection, the library pool was purified using AMPure PB beads. Size selection was performed a BluePippin instrument (Sage Science, Beverly, US) using BluePippin with dye free, 0.75% Agarose Cassettes and S1 Marker (Sage Science) wherein the selection cut-off was set at 6000 bp. Library pool concentration and size was again assessed using a Thermo Fisher Scientific Qubit 4.0 flurometer and an Advanced Analytical FEMTO Pulse instrument (as described above), respectively. PacBio Sequencing primer v4 and Sequel DNA Polymerase 3.0 were annealed and bound, respectively, to the DNA template libraries. The polymerase binding time was 1 h and the complex was cleaned using 1.2 X AMPure PB beads. The libraries were loaded at an on-plate concentration of 150pM using adaptive loading, along with the use of Spike-In internal control. SMRT sequencing was performed in CLR mode on the Sequel IIe with Sequel Sequencing kit 3.0, SMRT Cells 8M, a 2h pre-extension followed by a 15 h movie time and via PacBio SMRT Link v10.1. Thereafter, the CCS generation and barcode demultiplexing workflow was run in SMRT Link v10.1. All steps from gDNA extraction to sequencing data generation were performed at the Next Generation Sequencing Platform, University of Bern, Switzerland. The raw sequencing data is available from the European Nucleotide Archive (http://www.ebi.ac.uk/ena) with the study accession XYZ (sample IDs ABC to DEF).

#### Genome assembly

We utilized similar pipelines to assemble the genomes of all MRB strains. For set 1 (PacBio *and* Illumina sequence data), the fasta sequences of the ‘continuous long reads’ (CLRs), as extracted from the BAM files using samtools v. 1.10 (15), were used for assembly conducted with Flye v. 2.9 (16). Since these strains were also sequenced on Illumina sequencers, the CLR assembly was corrected with Illumina reads. The reads were first mapped to the assembly using the Burrows-Wheeler Aligner BWA, v 0.7.8 (17). The resulting SAM file was then sorted and indexed using samtools v. 1.10 before using Pilon v. 1.24 (18) to correct the assemblies.

For sets 2 to 4 (generated on Illumina sequencers), the raw, paired end fastq sequences were trimmed using fastp v. 0.20.1 (19) with default options. Read quality was assessed with fastQC v. 0.11.7 (7). These genomes were assembled using the SPAdes assembler v. 3.14.0 (20) with the options ‘--isolate –k 21,33,55,77,99,127 --cov-cutoff ‘auto’’. The quality of the assemblies was assessed with Quast v. 4.6.0 (21), BUSCO v. 5.1.3 (22) and checked for contamination with ConFindr v. 0.7.2 (23). The genomes were then annotated with the NCBI procaryotic genome assembly pipeline PGAP, v. 2022-04-14 (24). The annotated genomes were functionally annotated with EggNog v. 5.0.1 (25) and orthologue genes were determined using OrthoFinder v. 2.3.8 (26). The annotated assemblies were then integrated into a local instance of OpenGenomeBrowser (27) hosted at the Interfaculity Bioniformatics Unit (University of Bern). The genome assemblies and annotations have been deposited at BioProject database under accession numbers ABC to DEF (Dataset S1).

### Supplementary Figures

**Figure S1.**
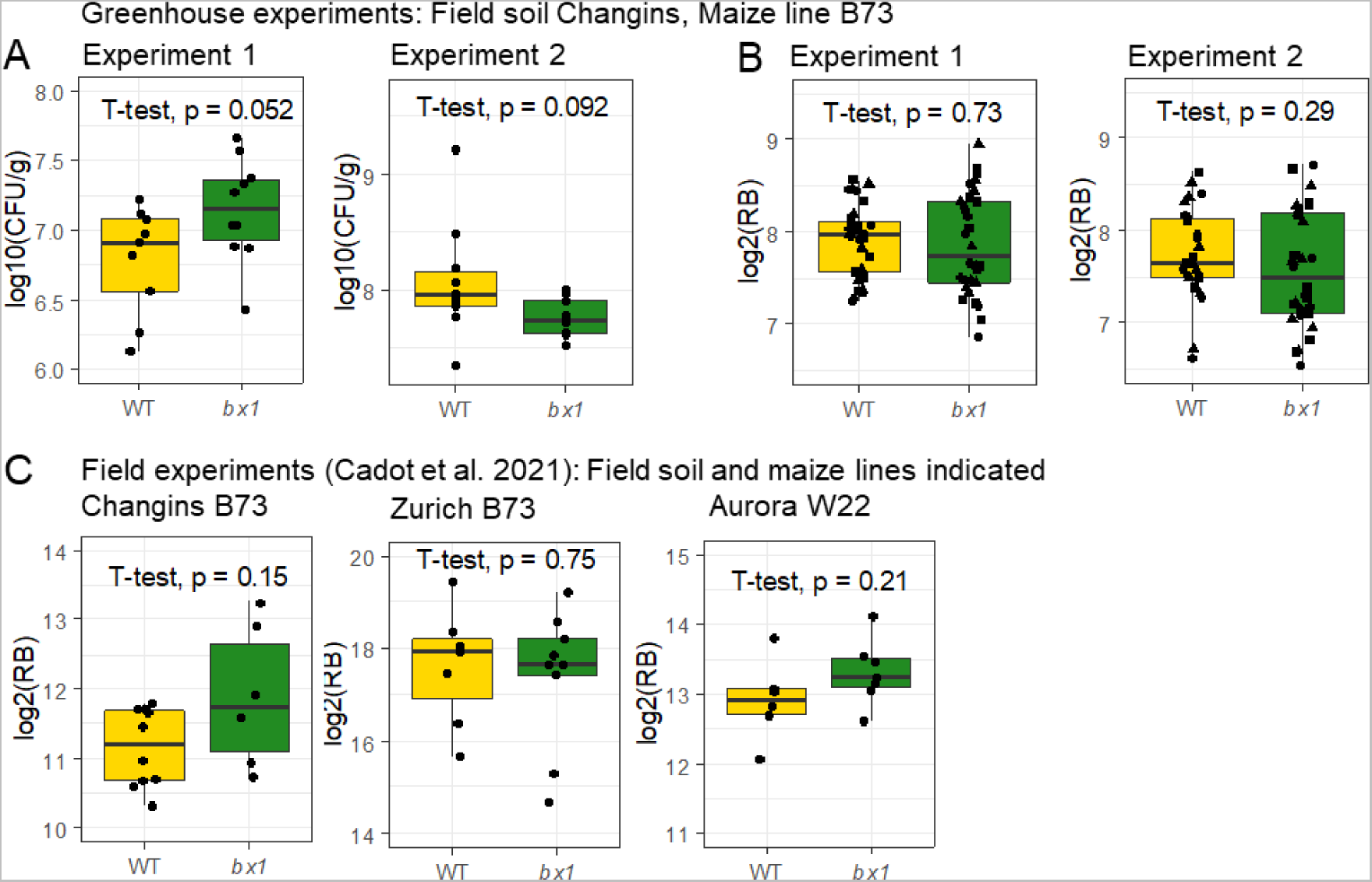
Bacterial community size on maize roots by microbiological and qPCR analyses. **A)** Bacterial root colonization was assessed by plating colony forming units (CFU) expressed as log10 CFU / g roots and tested for counts plotted, and statistically significant differences between wild-type and bx1 plants. **B)** DNA extracts from the same plants were used for qPCR analysis. The bacterial signal, derived from 16S rRNA primers 799F and 904R, was normalized relative to the plant signal of the plant actin gene (ZmActin1) expressed as log2(RB), RB = relative bacterial gene signal (Eplant gene ^Ct plant gene^/ E_16S_ ^Ct16S^) C) DNA extracts from maize roots grown in three field experiments published in Cadot et. al. 2021. Results from t-test between wild-type and bx1 are shown in the panel.

**Figure S2:**
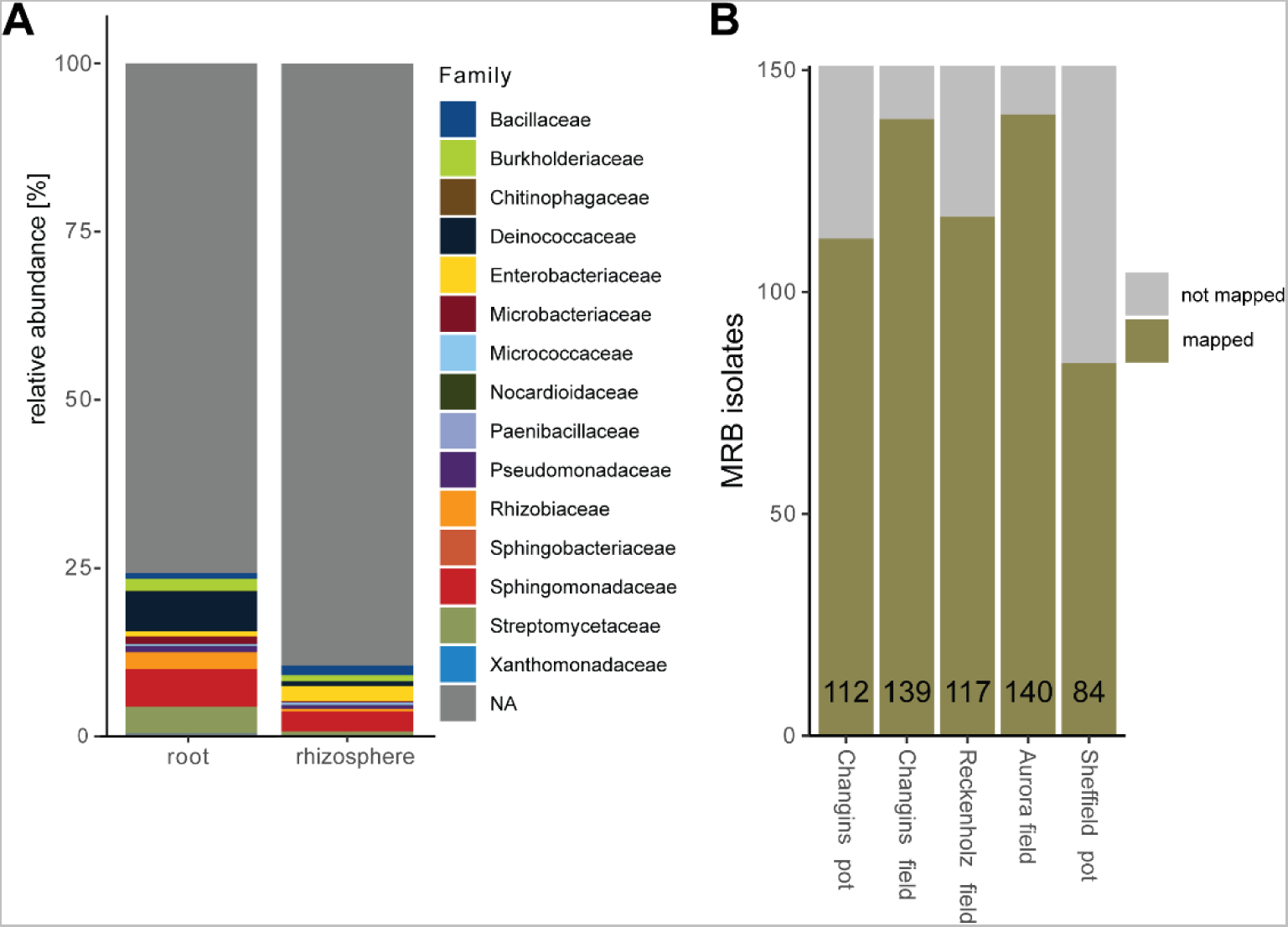
Mapping of MRB strains to maize root and rhizosphere microbiome datasets. **A)** Cumulative abundance of MRB strains, reported at family level, in the root and rhizosphere profiles of wild-type B73 maize plants, from which the MRB strains were isolated from. This was a greenhouse experiment with pots filled with natural field soil from the Changins site. The microbiome data corresponds to the feedback experiment reported in Hu et al. 2018. **B)** Number of MRB isolates mapping to abundant community members (> 0.1 % abundance) in root microbiome datasets of maize grown in greenhouse and field experiments (Changins field data from Hu et al. 2018; Reckenholz and Aurora data from Cadot et al. 2021) or a greenhouse experiment with field soil (Sheffield data from Cotton et al. 2019).

**Figure S3.**
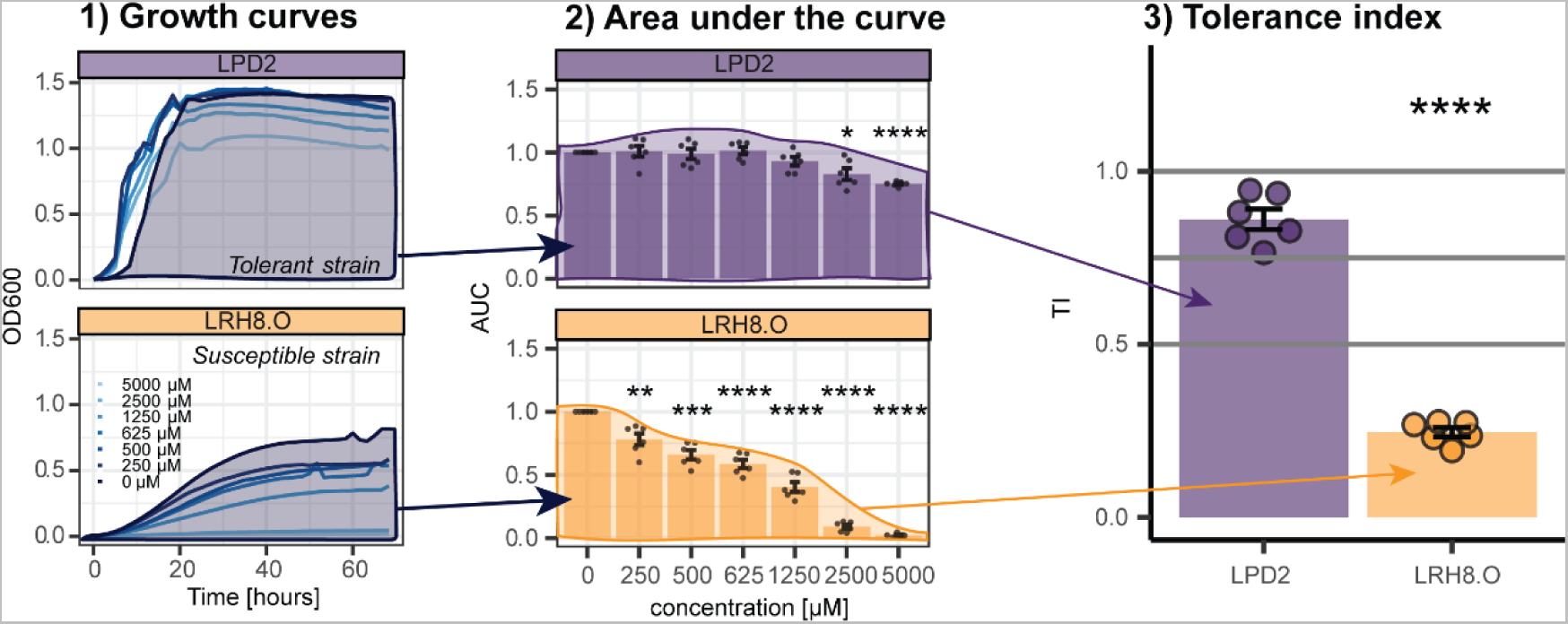
In vitro growth of maize root bacteria in MBOA. **A)** Bacterial growth curves (OD_600_) of a representative tolerant strain of Pseudomonadaceae (LPD2) and a representative susceptible strain of Rhizobiaceae (LRH8.O) at different concentrations of MBOA over a time course of 68 hours. **B)** Area under the curve (AUC), normalized to the BX-free control treatment **C)** Tolerance index (TI). Means ± SE bar graphs and individual datapoints are shown (n = 6). Results of pairwise t-test is shown inside the panels, p-value < 0.05 = *.

**Figure S4:**
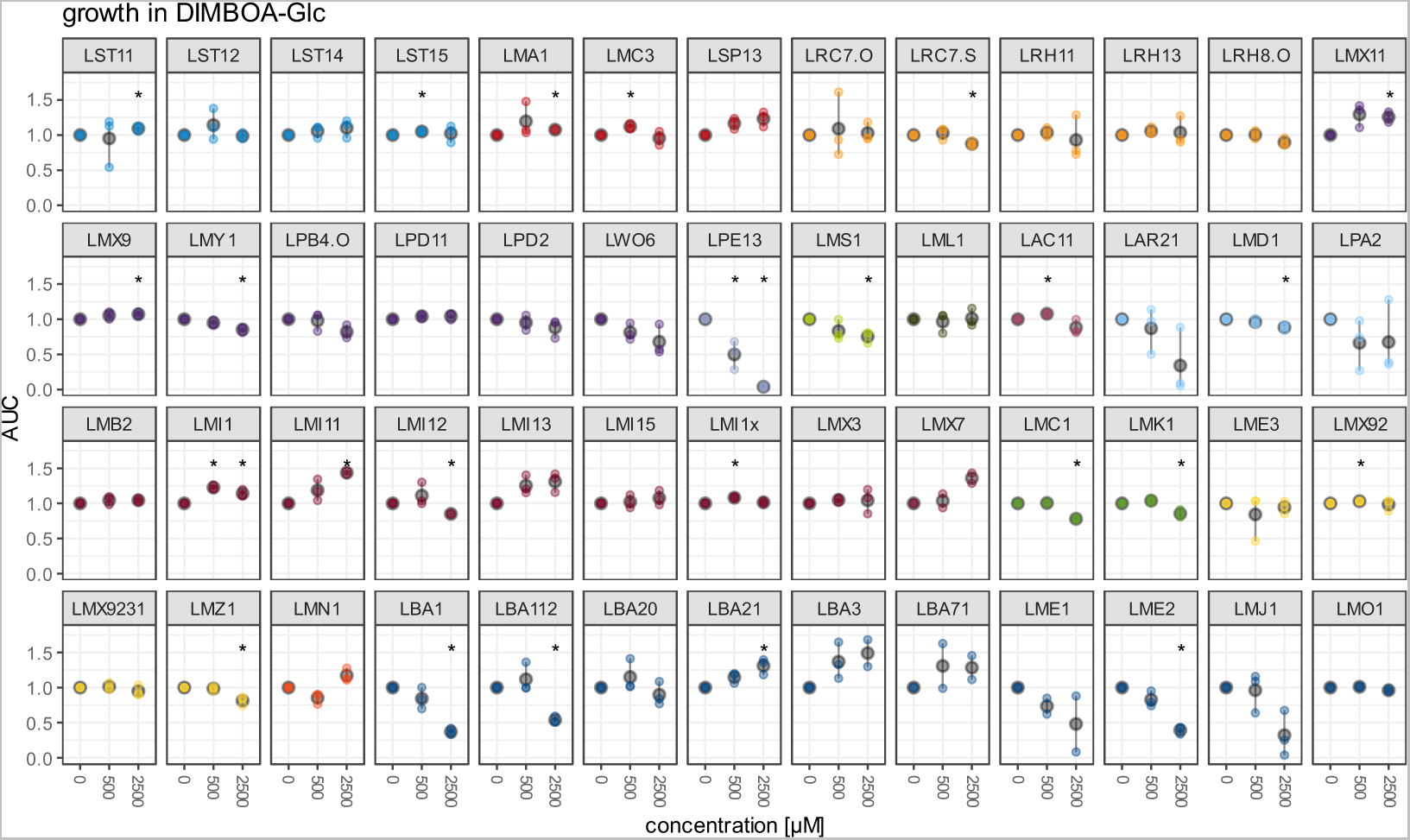
Growth of maize root bacteria in DIMBOA-Glc. AUC of maize root bacteria in 500 and 2500 µM of DIMBOA-Glc. Asterisks indicate significant differences (t-test) to the control; p-value < 0.05 = *.

**Figure S5:**
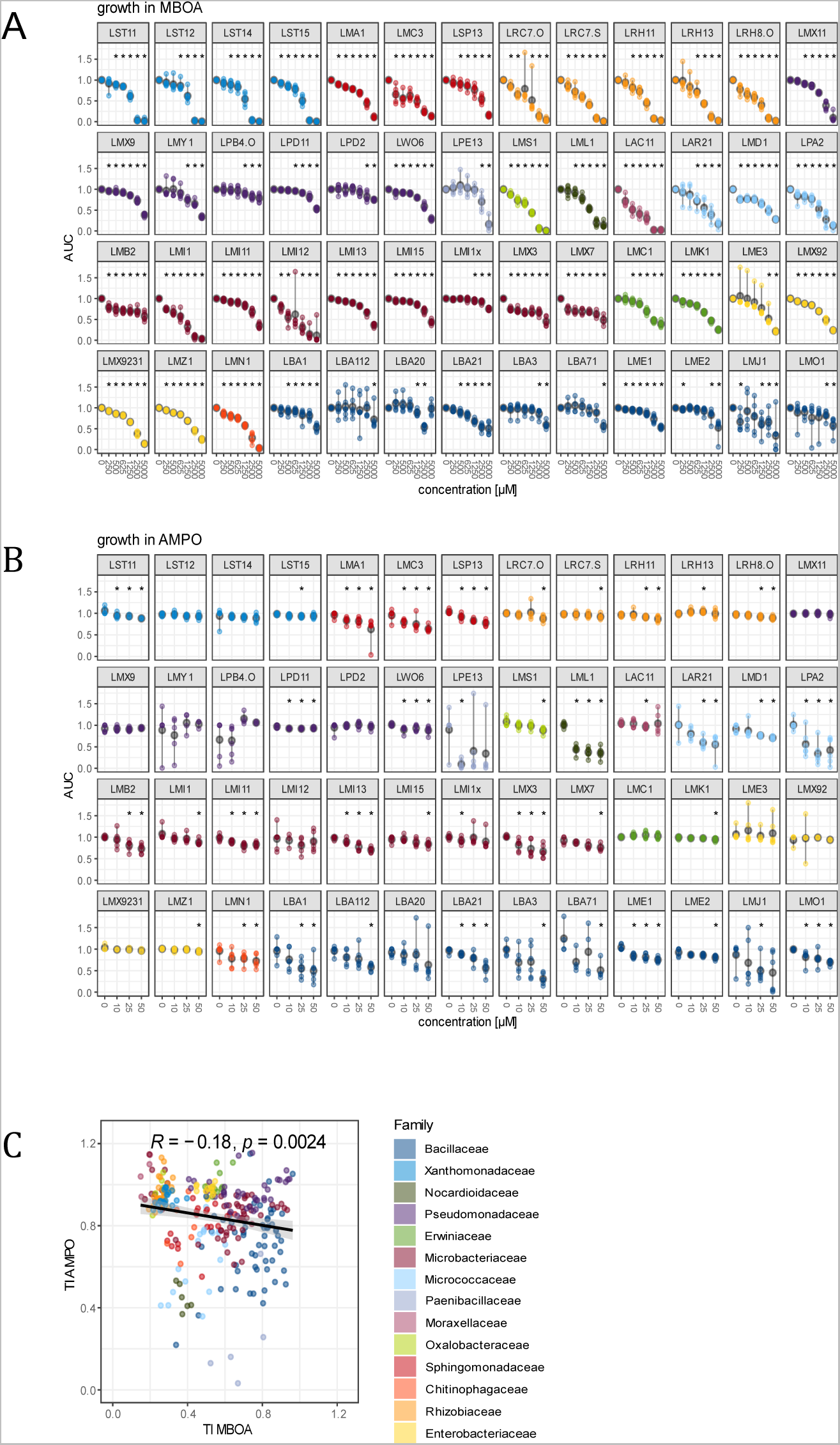
Growth of maize root bacteria in methoxylated benzoxazinoids and aminophenoxazinones. **A)** AUC of MRB strains in MBOA 50-5000 µM and **B)** AMPO 10-50 µM. Asterisks indicate significant differences (t-test) to the control; p-value < 0.05 = *. C) Correlation between TI of MBOA and AMPO. The correlation coefficient R and p-value of the Pearson’s product-moment correlation are shown on top.

**Figure S6:**
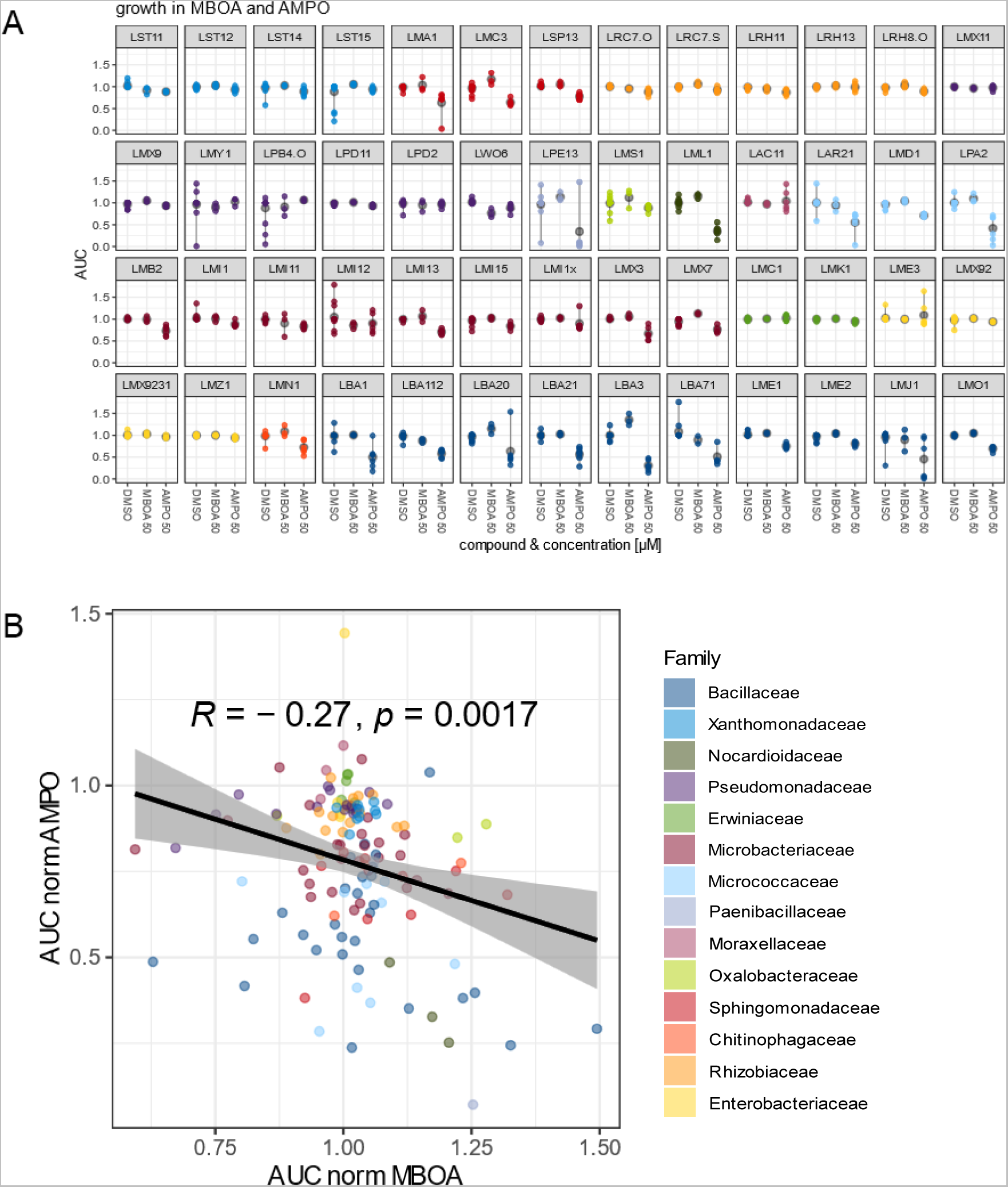
Growth of maize root bacteria in 50 µM of MBOA and AMPO. **A)** Comparison of AUC in control treatment (DMSO) with AUC in MBOA 50 µM and AMPO 50 µM. **B)** Correlation between AUC of MBOA and AMPO, each at 50 µM. The correlation coefficient R and p-value of the Pearson’s product-moment correlation are shown above.

**Figure S7.**
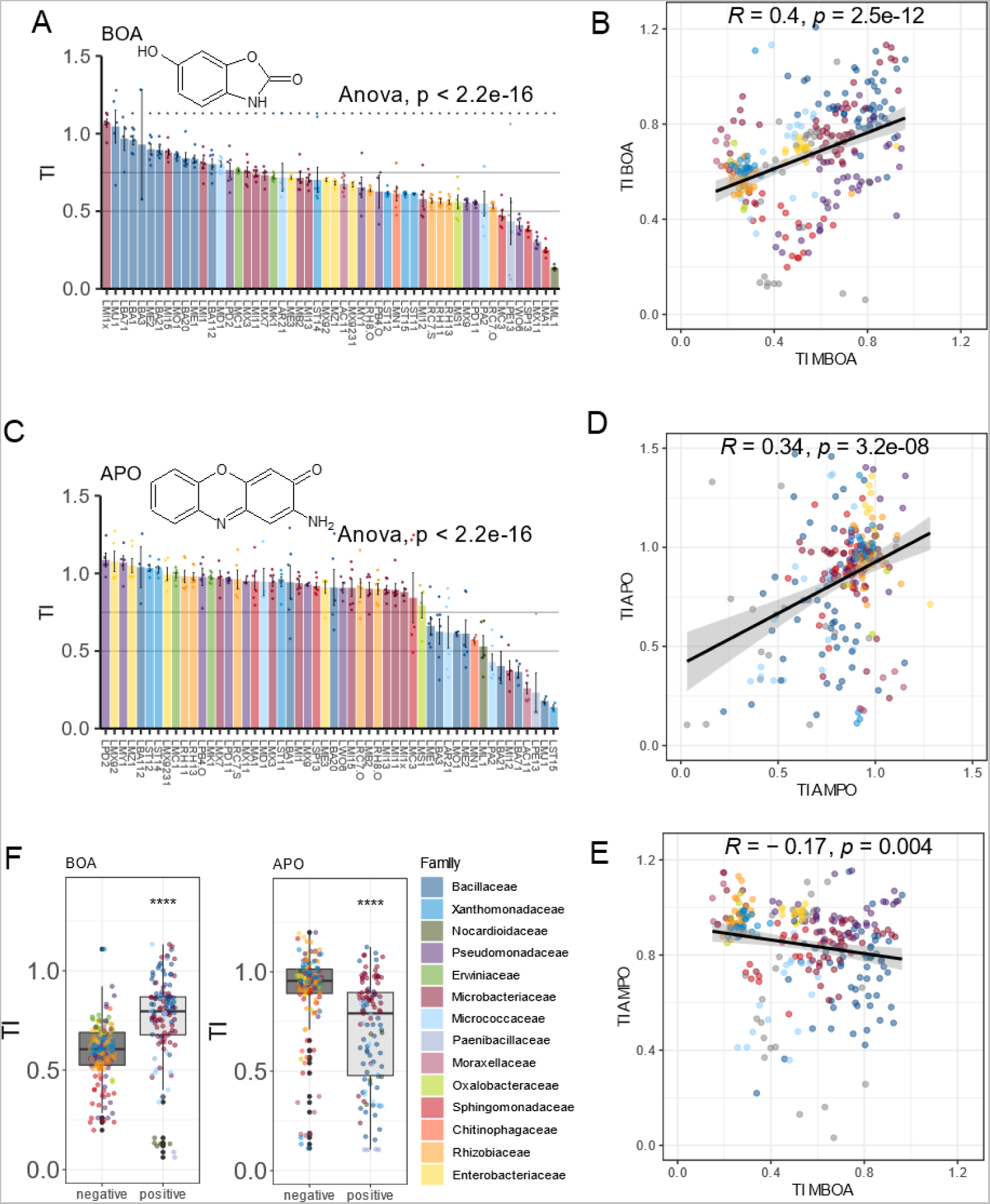
Tolerance of maize root bacteria to non-methoxylated benzoxazinoids. **A)** Bacterial strains are colored according to the family showing and their TI for BOA is reported. **B)** Correlation between TIs of BOA and MBOA. **C)** Bacterial strains are colored according to the family with their TI for APO. **D)** Correlation between TI of APO and AMPO. E) Correlation between TIs of APO and BOA. F) TI summarized for Gram-positive and Gram-negative MRB strains to BOA and APO. Graphs (A and C) report means ± SE with individual datapoints (n = 6 for BOA and APO) with statistic results of ANOVA are shown inside the panels, p-value < 0.05 = *. B, D and E) The correlation coefficient R and p-value of the Pearson’s product-moment correlation are shown on top.

**Figure S8:**
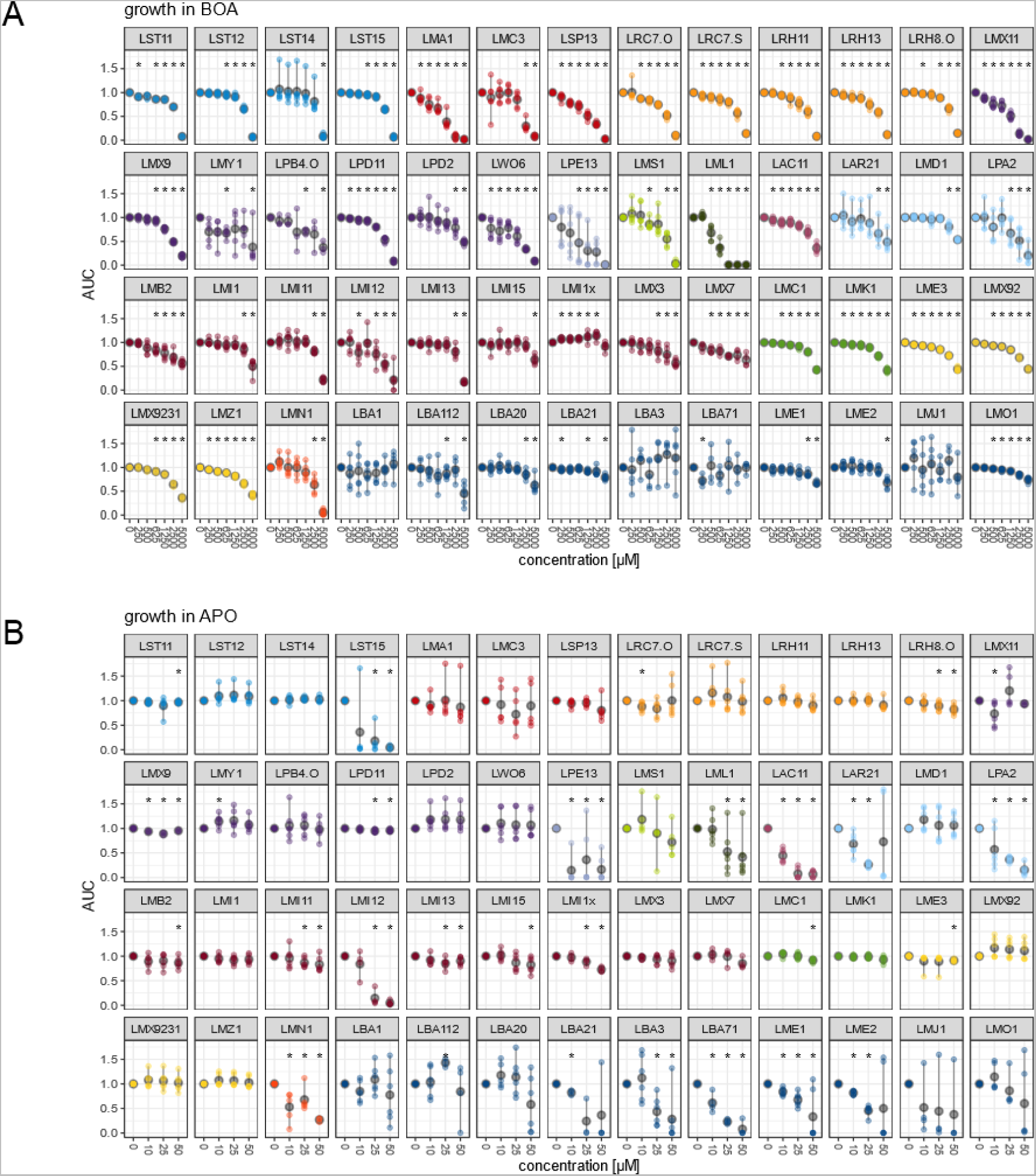
Growth of maize root bacteria in non-methoxylated benzoxazinoids and aminophenoxazinones. AUC of bacteria in **A)** in BOA 500-5000 µM, and **B)** APO 10-50 µM. Asterisks indicate significant differences (t-test) to the control; p-value < 0.05 = *.

**Figure S9:**
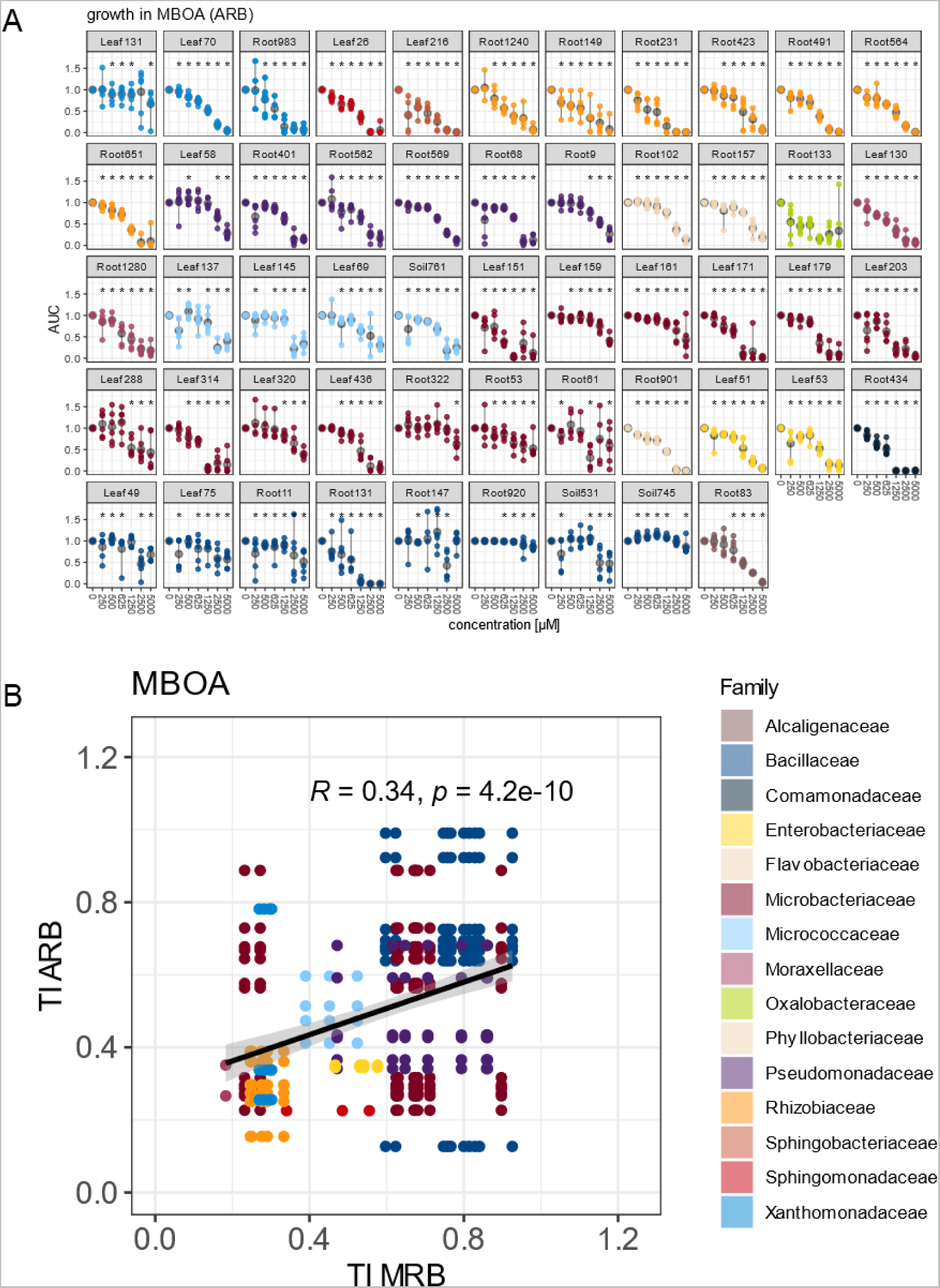
Growth of Arabidopsis root bacteria in MBOA. **A)** AUC of AtSphere bacteria in MBOA 250-5000 μM. **B)** Correlation between TI of MRB and ARB in MBOA.

**Figure S10:**
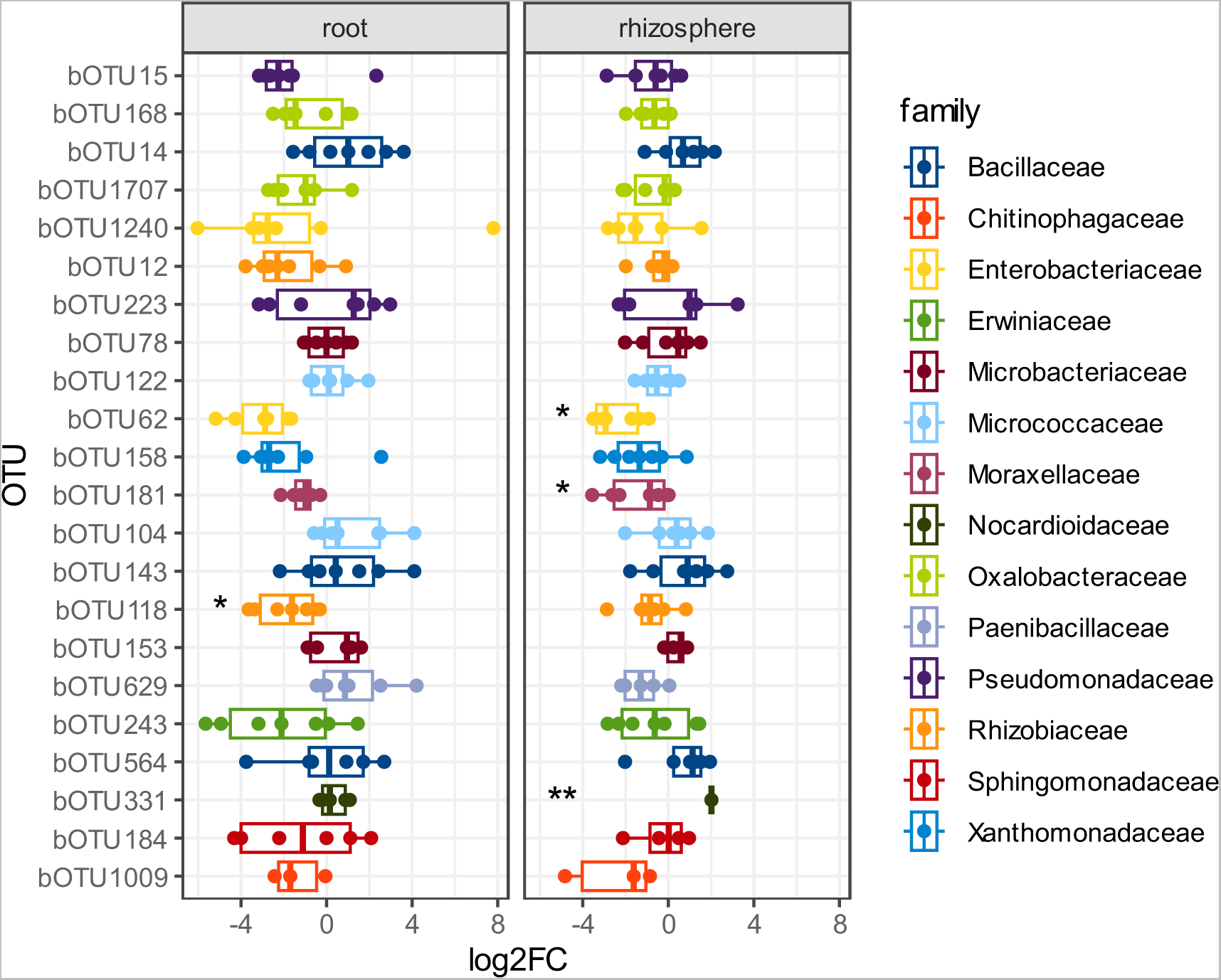
Differential abundance of OTUs, to which the MRB strains map, on roots and in rhizosphere profiles of field-grown maize. MRB isolates were mapped to the operational taxonomic units (OTUs) of the microbiome dataset of Hu et al. 2018, where wild-type and bx1 mutant lines were grown in a field experiment in Changins. log2fold changes > 1 indicate OTU enrichment on wild-type while values < 1 indicate OTU depletion on wild-type plants. Asterisks indicate significant differences of absolute abundances of the OTU on both genotypes. These values were used to calculate log2fold ratio (t-test)

**Figure S11:**
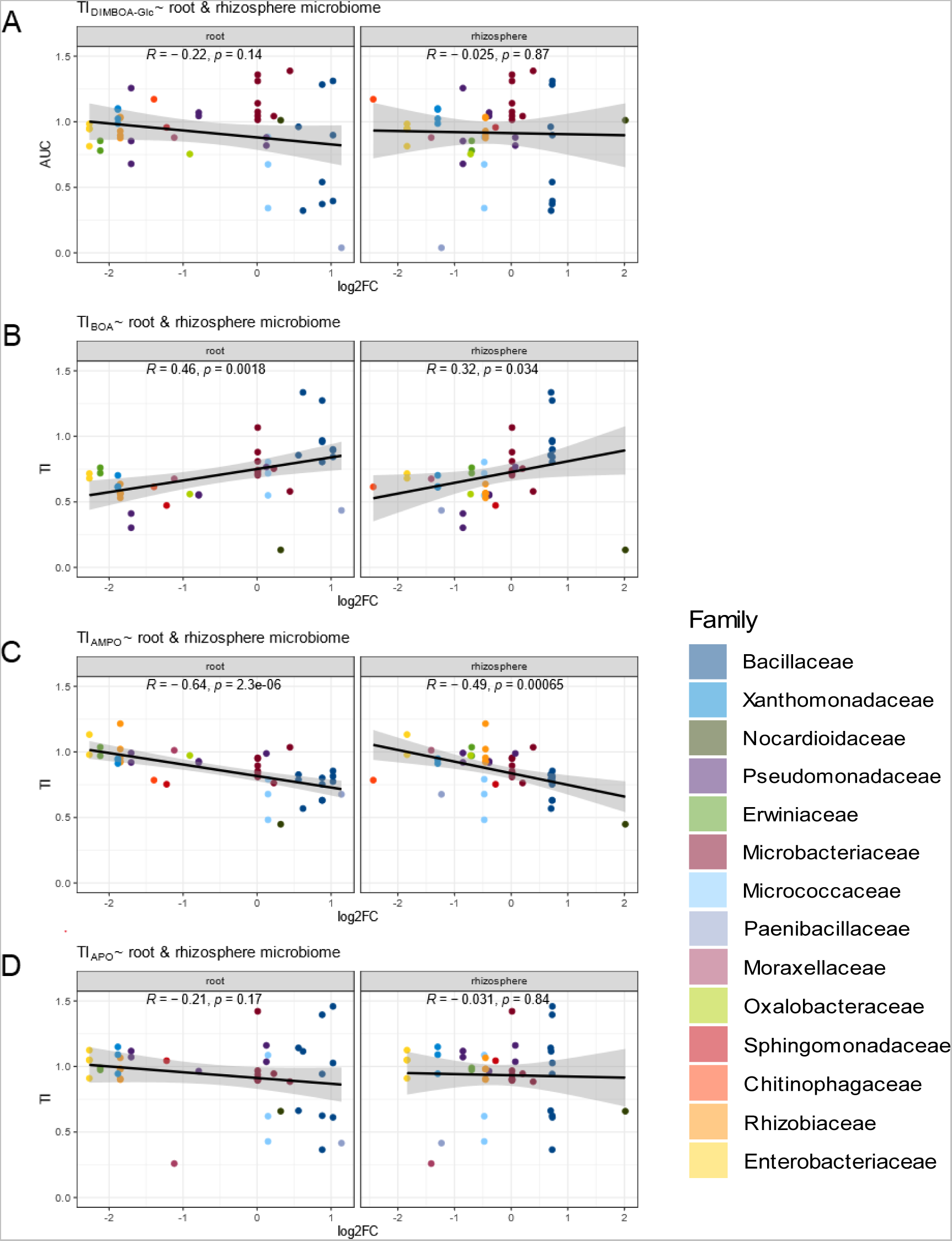
Correlation of bacterial BX tolerance with BX-dependent abundance on maize roots. **A)** Correlation between log2fold change (log2FC) of the respective OTUs in the field on wild-type to bx1 roots and in rhizosphere with AUC DIMBOA-Glc, **B)** with TI AMPO, **C)** with TI BOA and **D)** with TI APO. Pearson’s product-moment correlation test was performed. The correlation coefficient R and p-value of the Pearson’s product-moment correlation are shown on top. p-value < 0.0001 = ****, < 0.001 = ***, < 0.01 = **, 0.05 = *

### Supplementary Tables

**Table S1:**
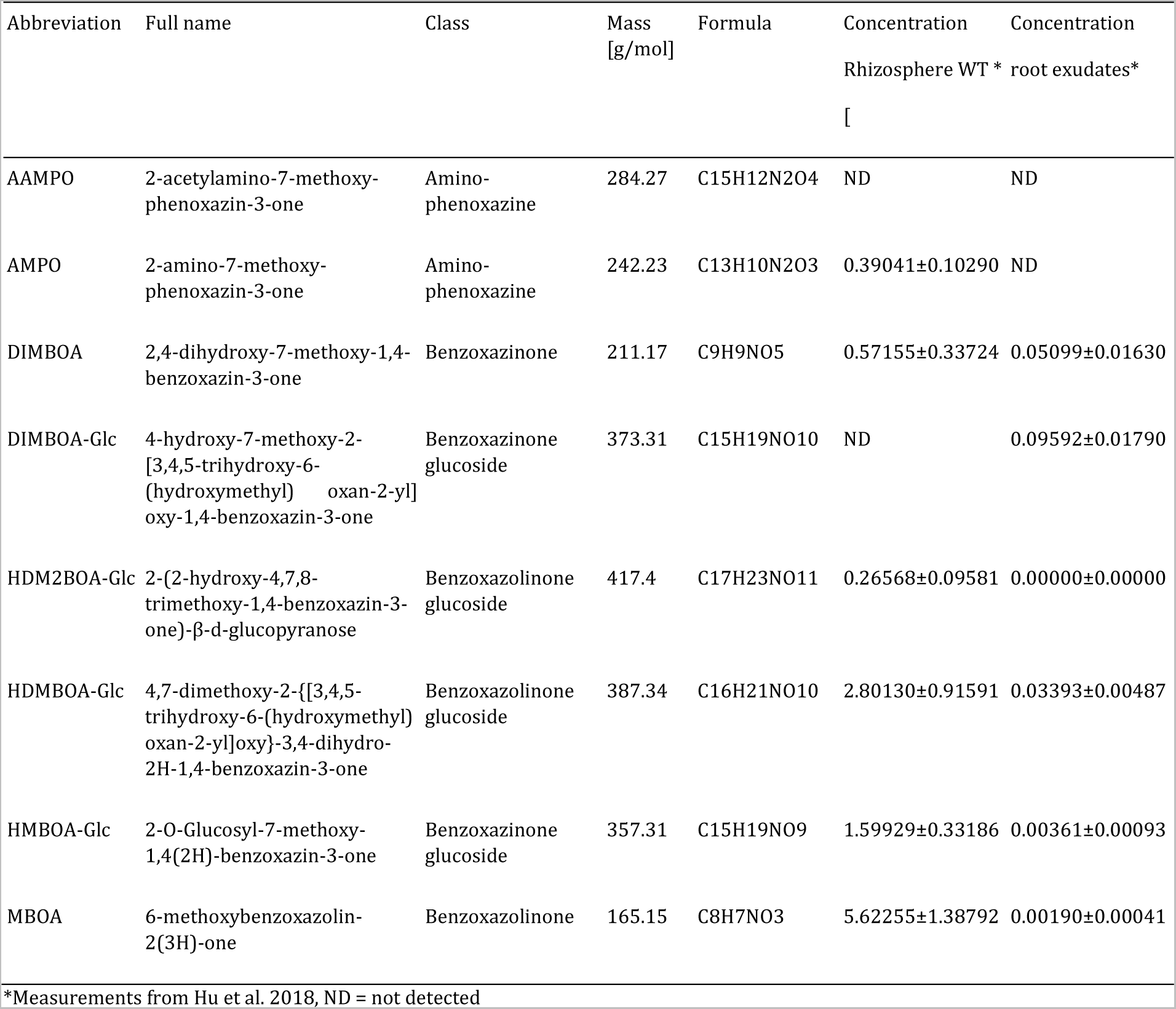
Abbreviation benzoxazinoid metabolites

| Abbreviation | Full name | Class | Mass<br>[g/mol] | Formula | Concentration<br>Rhizosphere WT * | Concentration<br>root exudates* |
| --- | --- | --- | --- | --- | --- | --- |
| AAMPO | 2-acetylamino-7-methoxy-phenoxazin-3-one | Amino-phenoxazine | 284.27 | C15H12N2O4 | ND | ND |
| AMPO | 2-amino-7-methoxy-phenoxazin-3-one | Amino-phenoxazine | 242.23 | C13H10N2O3 | 0.39041±0.10290 | ND |
| DIMBOA | 2,4-dihydroxy-7-methoxy-1,4-benzoxazin-3-one | Benzoxazinone | 211.17 | C9H9NO5 | 0.57155±0.33724 | 0.05099±0.01630 |
| DIMBOA-Glc | 4-hydroxy-7-methoxy-2-[3,4,5-trihydroxy-6-(hydroxymethyl)oxan-2-yl]oxy-1,4-benzoxazin-3-one | Benzoxazinone glucoside | 373.31 | C15H19NO10 | ND | 0.09592±0.01790 |
| HDM2BOA-Glc | 2-(2-hydroxy-4,7,8-trimethoxy-1,4-benzoxazin-3-one)-β-d-glucopyranose | Benzoxazolinone glucoside | 417.4 | C17H23NO11 | 0.26568±0.09581 | 0.00000±0.00000 |
| HDMBOA-Glc | 4,7-dimethoxy-2-[[3,4,5-trihydroxy-6-(hydroxymethyl)oxan-2-yl]oxy]-3,4-dihydro-2H-1,4-benzoxazin-3-one | Benzoxazolinone glucoside | 387.34 | C16H21NO10 | 2.80130±0.91591 | 0.03393±0.00487 |
| HMBOA-Glc | 2-O-Glucosyl-7-methoxy-1,4(2H)-benzoxazin-3-one | Benzoxazinone glucoside | 357.31 | C15H19NO9 | 1.59929±0.33186 | 0.00361±0.00093 |
| MBOA | 6-methoxybenzoxazolin-2(3H)-one | Benzoxazolinone | 165.15 | C8H7NO3 | 5.62255±1.38792 | 0.00190±0.00041 |

\*Measurements from Hu et al. 2018, ND = not detected

**Table S2.**
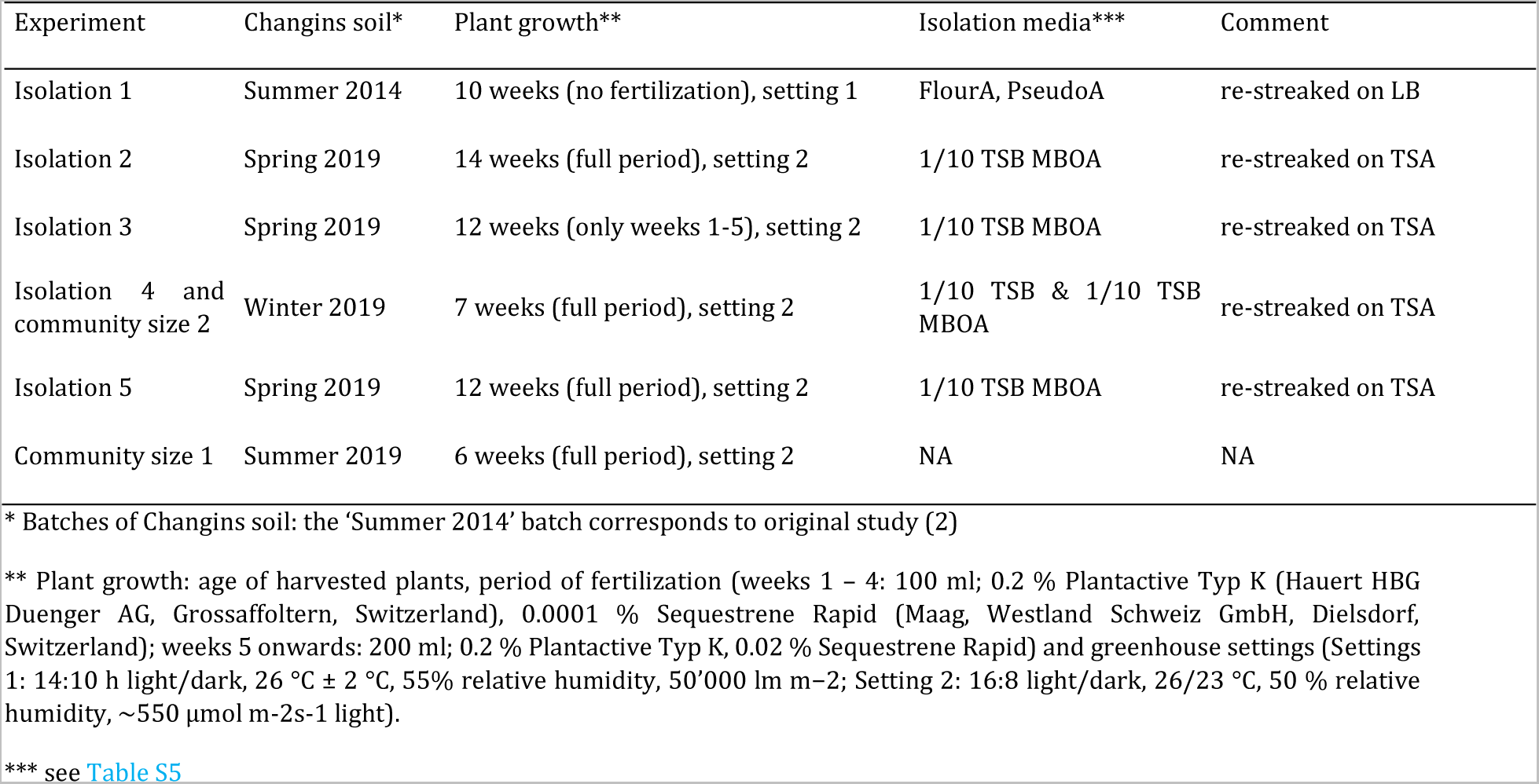
Experiments and plant growth conditions

| Experiment | Changins soil* | Plant growth** | Isolation media*** | Comment |
| --- | --- | --- | --- | --- |
| Isolation 1 | Summer 2014 | 10 weeks (no fertilization), setting 1 | FlourA, PseudoA | re-streaked on LB |
| Isolation 2 | Spring 2019 | 14 weeks (full period), setting 2 | 1/10 TSB MBOA | re-streaked on TSA |
| Isolation 3 | Spring 2019 | 12 weeks (only weeks 1-5), setting 2 | 1/10 TSB MBOA | re-streaked on TSA |
| Isolation 4 and community size 2 | Winter 2019 | 7 weeks (full period), setting 2 | 1/10 TSB & 1/10 TSB MBOA | re-streaked on TSA |
| Isolation 5 | Spring 2019 | 12 weeks (full period), setting 2 | 1/10 TSB MBOA | re-streaked on TSA |
| Community size 1 | Summer 2019 | 6 weeks (full period), setting 2 | NA | NA |

\* Batches of Changins soil: the 'Summer 2014' batch corresponds to original study (2)
\*\* Plant growth: age of harvested plants, period of fertilization (weeks 1 – 4: 100 ml; 0.2 % Plantactive Typ K (Hauert HBG Duenger AG, Grossaffoltern, Switzerland), 0.0001 % Sequestrene Rapid (Maag, Westland Schweiz GmbH, Dielsdorf, Switzerland); weeks 5 onwards: 200 ml; 0.2 % Plantactive Typ K, 0.02 % Sequestrene Rapid) and greenhouse settings (Settings 1: 14:10 h light/dark, 26 °C ± 2 °C, 55% relative humidity, 50'000 lm m<sup>-2</sup>; Setting 2: 16:8 light/dark, 26/23 °C, 50 % relative humidity, ~550 µmol m<sup>-2</sup>s<sup>-1</sup> light).
\*\*\* see Table S5

**Table S3.**
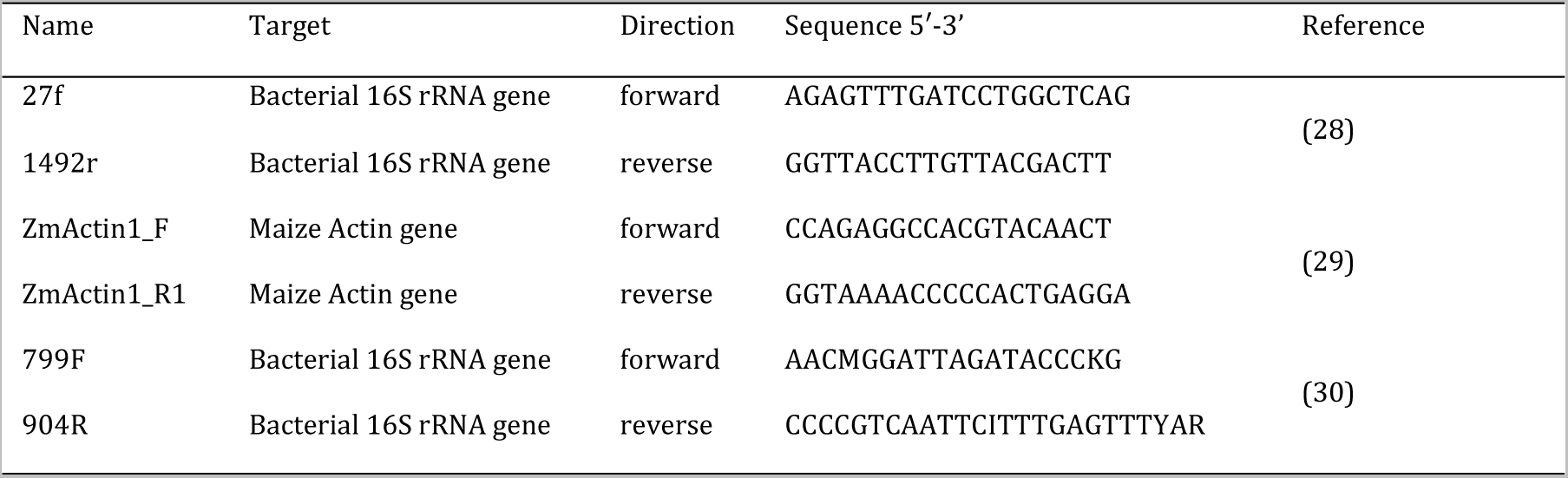
PCR primer sequences

**Table S4.**
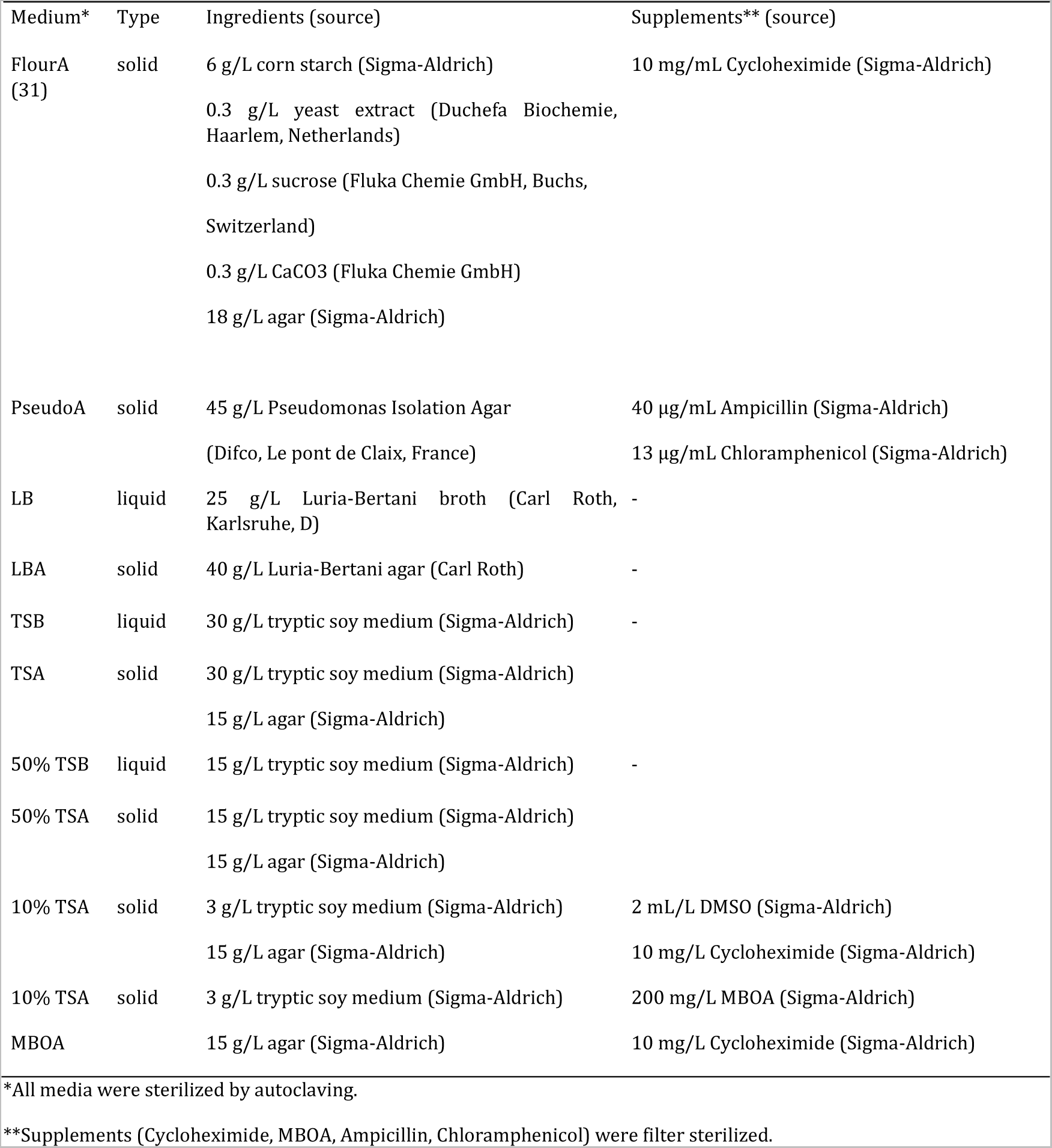
Media uses for isolation of maize root bacteria

**Table S5.** Stock solutions of compounds used for in vitro growth assays

| Compound | mol. weight | stock conc. [mM] | stock mg/mL | solvent |
| --- | --- | --- | --- | --- |
| DIMBOA-Glc | 373.1 | 500 | 186.55 | DMSO |
| MBOA | 165 | 606 | 100 | DMSO |
| BOA | 135.1 | 500 | 67.55 | DMSO |
| AMPO | 242.23 | 15 | 3.6 | DMSO |
| APO | 212.21 | 15 | 3.18 | DMSO |
| Ctrl | 0 | 0 | 0 | DMSO |

### Supplementary Datasets

**Dataset S1. MRB strain collection sequences.** This table contains detailed information about the taxonomic assignment of the MRB isolates. Further information on the isolation experiment, the plant, the extract, and the isolation media are included. The partial sequence of the 16S rRNA gene obtained by Sanger sequencing along with the primer used is listed. For each strain it is indicated if and with which method the genome was sequenced.

**Dataset S2. MRB strain collection mapping.** The mapping of MRB isolates to the microbiome profiles of the maize roots, where they were isolated from (pot experiment with Changins soil) indicating the identity to the taxonomic units.

